# Demystifying the Visual Word Form Area: Visual and Nonvisual Response Properties of Ventral Temporal Cortex with precision fMRI

**DOI:** 10.1101/2023.06.15.544824

**Authors:** Jin Li, Kelly J. Hiersche, Zeynep M. Saygin

## Abstract

The functional nature of the visual word form area (VWFA), a region in left ventrotemporal cortex (VTC), remains debated after over two decades of investigation. Here we use precision fMRI to comprehensively examine the VWFA’s responses to numerous visual and nonvisual stimuli. We find that the VWFA is a distinct region that while showing moderate activation to non-word visual stimuli, its response to words towers above other conditions. Interestingly, the VWFA is also the only VTC region uniquely engaged in auditory language processing, in contrast with the ubiquitous attentional effect within the VTC. However, this language selectivity is dwarfed by its visual responses. This suggests the VWFA primarily functions as a visual look-up orthographic dictionary. Searching within the entire VTC further reveals an additional language cluster in anterior fusiform preferring general visual semantics. Our detailed examination of the VWFA in direct comparison to other VTC regions clarifies the long-standing controversies about its functionality.

## Introduction

The ventrotemporal cortex (VTC) consists of numerous regions that each specialize in perceiving abstract visual stimulus categories (e.g., faces, objects, bodies, and places). The visual word form area (VWFA) is perhaps one of the most fascinating of these VTC regions because it is specialized for processing a recent human invention: reading (Cohen et al., 2002; McCandliss et al., 2003). This functional specialization, as well as the experience-dependent nature of the VWFA <u>(e.g., Baker et al., 2007; Dehaene et al., 2015)</u>, make it a prime example for understanding the functional organization of the human brain. However, there is still debate over whether the VWFA is specialized *specifically* for visual words, which precludes researchers from digging deeper into the functional characteristics of the VWFA and how the human brain has the capacity to dedicate cortical tissue for new symbolic representations.

The key argument against the idea of a region that is dedicated to visual words is that the VWFA is also activated for other meaningful, non-word stimuli (e.g., Price & Devlin, 2003, 2011). Proponents of this view argued that given the relatively recent invention of written script, the response to visual words is likely repurposed from other functionally specialized regions (e.g., Dehaene & Cohen, 2007), and still maintains other functions (Vogel et al., 2014). Studies that supported this view have shown that responses to words in this region were not qualitatively higher than responses to line-drawings of objects, false fonts (Ben-Shachar et al., 2007) or unfamiliar scripts (Xue & Poldrack, 2007). Therefore, some argued that the anatomical location of the VWFA, the posterior fusiform gyrus, is involved in complex shape processing (Roberts et al., 2013) more generally (Mei et al., 2010; Neudorf et al., 2022). However, we argue that the VWFA’s responses to non-word stimuli in isolation (i.e. not in comparison to its responses to word stimuli) should not be taken as evidence against the VWFA’s word selectivity. In fact, even the fusiform face area responds to non-face stimuli (e.g., Kanwisher et al., 1997). Instead, to better probe the function of the VWFA, one should ask 1) whether the VWFA shows similar functional characteristics as other category-selective VTC regions that respond significantly (and robustly, e.g., ∼twice as much) higher to the words than non-word categories (Kay & Yeatman, 2017), and 2) what is the functional profile of the VWFA in terms of its preferences to non-word categories.

Another argument against the VWFA’s specialization for visual words is its involvement in auditory language processing (e.g., Ludersdorfer et al., 2016; Planton et al., 2019; Price & Devlin, 2003). In congenitally blind individuals, the site of the VWFA responds to both Braille words and auditory words but not tactile patterns or backward speech (Kim et al., 2017). Similarly, activation to auditory words was also found in sighted individuals (Ludersdorfer et al., 2016), and when participants are asked to selectively attend to speech via a rhyme judgment task, both frontotemporal language regions *and* the VWFA showed increased activation as compared to when melody was presented (Yoncheva et al., 2010). However, the FFA is also activated during imagery (e.g., O’Craven & Kanwisher, 2000) and Haptic stimuli of the faces (e.g., Kitada et al., 2009) and this does not mean that it is not specialized for faces. Instead, we should ask what is the functional nature of these activations to non-orthographic. Specifically, how do these auditory language responses compare to those for written language? And how is the VWFA uniquely involved in processing auditory language, as compared to adjacent VTC regions or the entirety of the VTC? Is the VWFA another node of the core language network that responds selectively to high-level linguistics like semantics and syntax, regardless of the input modality? For example, in addition to Braille and auditory words, the VWFA was also sensitive to grammatical complexity manipulation of auditory sentences (Kim et al., 2017). Alternatively, perhaps the VWFA mainly serves as a visual lookup dictionary for orthographic stimuli, which further passes visual inputs to frontotemporal language regions via its privileged connectivity with the language cortex.

Finally, the exact location and definition of the VWFA are not consistent in previous studies, which further makes it difficult to reach any consensus among studies regarding the function of the VWFA. Although located in approximately the same location across individuals, the VWFA is a small region, and the exact location varies from person to person (Glezer & Riesenhuber, 2013). However, a various previous studies examining the function of the VWFA relied on group activation maps or used anatomical coordinates (on a template brain) (Chen et al., 2019; Vogel, Miezin, et al., 2012; Vogel, Petersen, et al., 2012). These methods may not capture word-selective voxels because they do not account for individual variability. Further, previous studies investigating the VWFA usually solely defined the VWFA, or in addition to just one other VTC region as comparison. However, the mosaic-like organization of the VTC encompasses multiple category-selective regions that are located closely to each other and to the VWFA. Finally, a last point of inconsistency across previous studies was the control conditions used: the VWFA was initially defined using fixation/rest or checkboard stimuli (Cohen et al., 2000, 2002), or otherwise poorly controlled for visual complexity and general semantic processing. However, despite these limitations, subsequent studies continued referencing this anatomical location as the VWFA. Consequently, this lack of functional specificity in its initial definition could be a contributing factor to studies reporting activations in the VWFA during non-word processing tasks.

In the present study, rather than offering simple yes or no answers to the lingering debates about the VWFA, our goal is to systematically examine its functional characteristics using a wide range of visual and nonvisual stimuli. Specifically, we utilized precision fMRI to measure the subject-specific VWFA’s functional response profile across four distinct tasks spanning multiple sessions and encompassing a total of 13 experimental conditions. This allowed us to thoroughly probe the function of the VWFA in comparison with functionally related or spatially proximate regions. We assessed the VWFA’s activation in response to both high-level visual conditions and auditory language. The results not only demonstrate the robustness of the VWFA’s word selectivity but also provide insight into its activation during non-word processing, revealing its distinctive involvement in auditory language processing. By transcending the binary question of whether the VWFA exclusively processes words, our findings shed light on a more detailed picture of the VWFA’s responses. This understanding could potentially yield fresh insight into the development of the human brain’s functional organization.

## Results

### VWFA is selective to visual words, showing a distinct neural response signature from adjacent VTC regions

We first examined the functional response profile of the VWFA to a wide range of visual stimuli, as compared to other high-level category-selective regions (functional regions of interest (fROI), defined by contrasting the condition of interests with the remaining conditions in a localizer task; see **Methods**) within the left VTC. The main results focus on the left VTC given the left-lateralized nature of the VWFA (right VTC results in Supplementary **Figures S2 & S3** and **Table S2**). **Figure 1A** shows the fROIs in one example subject. Note that these spatially proximate regions make up the landscape of the VTC, with variability in the exact location of the fROIs across participants (all subjects **Figure S1),** highlighting the importance of defining subject-specific fROIs for multiple category-selective regions simultaneously to avoid blurring the boundaries of these regions in group analysis. We found that the VWFA responded significantly higher to visual words than all other conditions (paired samples t-tests, all p<0.001, **Table 1**, **Figure 1B**). When calculating the averaged time-course across the experimental block, response to words was higher than all other static and dynamic visual conditions (**Figure 1C**): the VWFA showed no preference for conditions other than words for the duration of the block.

**Figure 1.**
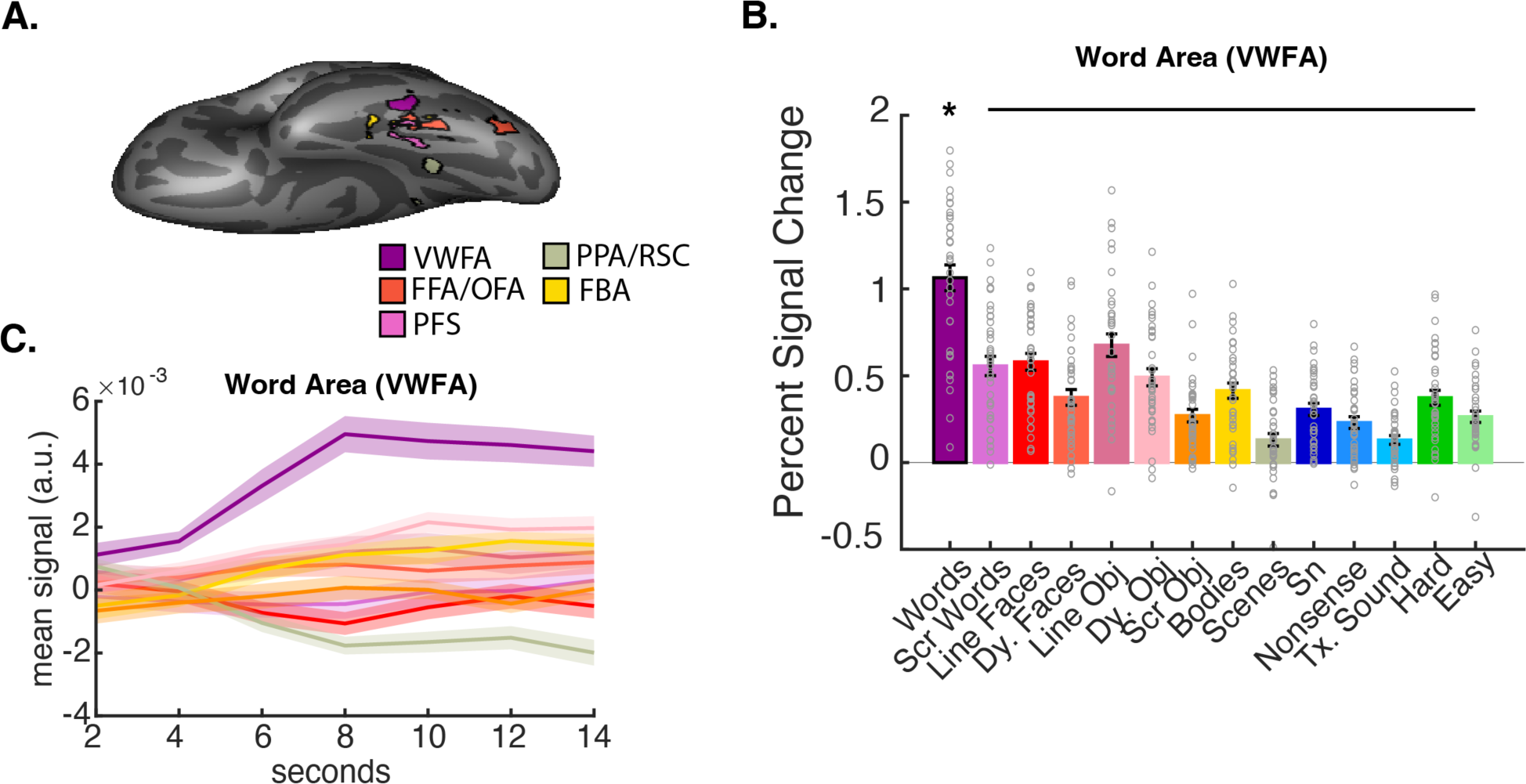
A. VTC fROIs in the left hemisphere for an example subject. B. Functional profile of the left VWFA. The mean percent signal change (with standard error bars) to various visual and non-visual conditions are plotted. The preferred category, words, has a thick black outline. Individual subject PSCs are shown with grey hollow circles. Significance is noted (*p<0.05, corrected for 13 total comparisons with Bonferroni-holm method) for words only, with a black line showing all conditions significantly lower than words. C. Averaged time-course of VWFA’s responses to blocks of different experimental conditions. Dark line for mean across all subjects and shading for standard error. VWFA, visual word form area; FFA, fusiform face area; OFA, occipital face area; PFS, posterior fusiform sulcus; PPA, parahippocampal place area; RSC, Retrosplenial Cortex (RSC); FBA, fusiform body area.

**Table 1.** Comparing the VWFA’s response to words with all other visual conditions (see Table S1 for other regions). **Bonferroni-Holm p<0.05.

| VWFA Words vs. | t (df) | Corrected p |
| --- | --- | --- |
| Scrambled Words | 7.03 (35) | $7.10 \times 10^{-8}$ ** |
| Line Faces | 7.93 (35) | $1.25 \times 10^{-8}$ ** |
| Dynamic Faces | 8.73 (35) | $1.56 \times 10^{-9}$ ** |
| Line Objects | 5.57 (35) | $2.87 \times 10^{-6}$ ** |
| Dynamic Objects | 7.19 (35) | $6.45 \times 10^{-8}$ ** |
| Scrambled Objects | 9.29 (35) | $3.95 \times 10^{-10}$ ** |
| Dynamic Bodies | 7.91 (35) | $1.25 \times 10^{-8}$ ** |
| Dynamic Scenes | 10.27 (35) | $3.36 \times 10^{-11}$ ** |

To what extent is the VWFA unique in its word-selective response? No other region showed the highest responses to visual words; instead, as expected, face-, object- and scene-selective regions showed significantly highest response for their preferred condition while the FBA did not show a clear categorical preference (**Figure 2, Table S1**. To further compare word-selective responses among VTC regions, we calculated the category selectivity index for each VTC region: the difference between PSC to preferred vs. non-preferred conditions (see **Methods** for details). A one-way rmANOVA comparing word selectivity among lVTC fROIs showed a significant main effect for fROI, (F(6,186)=33.89, p=1.81x10^-27^) (**Figure 3A**). Post-hoc comparison indicated that the VWFA is significantly more selective to words than all other fROIs (VWFA vs. all other fROIs, t(31)>5.13, p<1.47x10^-5^). Similarly, all other VTC fROIs showed significantly higher selectivity to their preferred categories vs. other adjacent fROIs (**Figure 3B**, **Table S3**), suggesting the functional distinctiveness of the VWFA as well as other VTC regions. Therefore, when localized with functional and spatial specificity, we confirmed that the VWFA is a unique VTC region that shows the strongest activation to written words.

**Figure 2.**
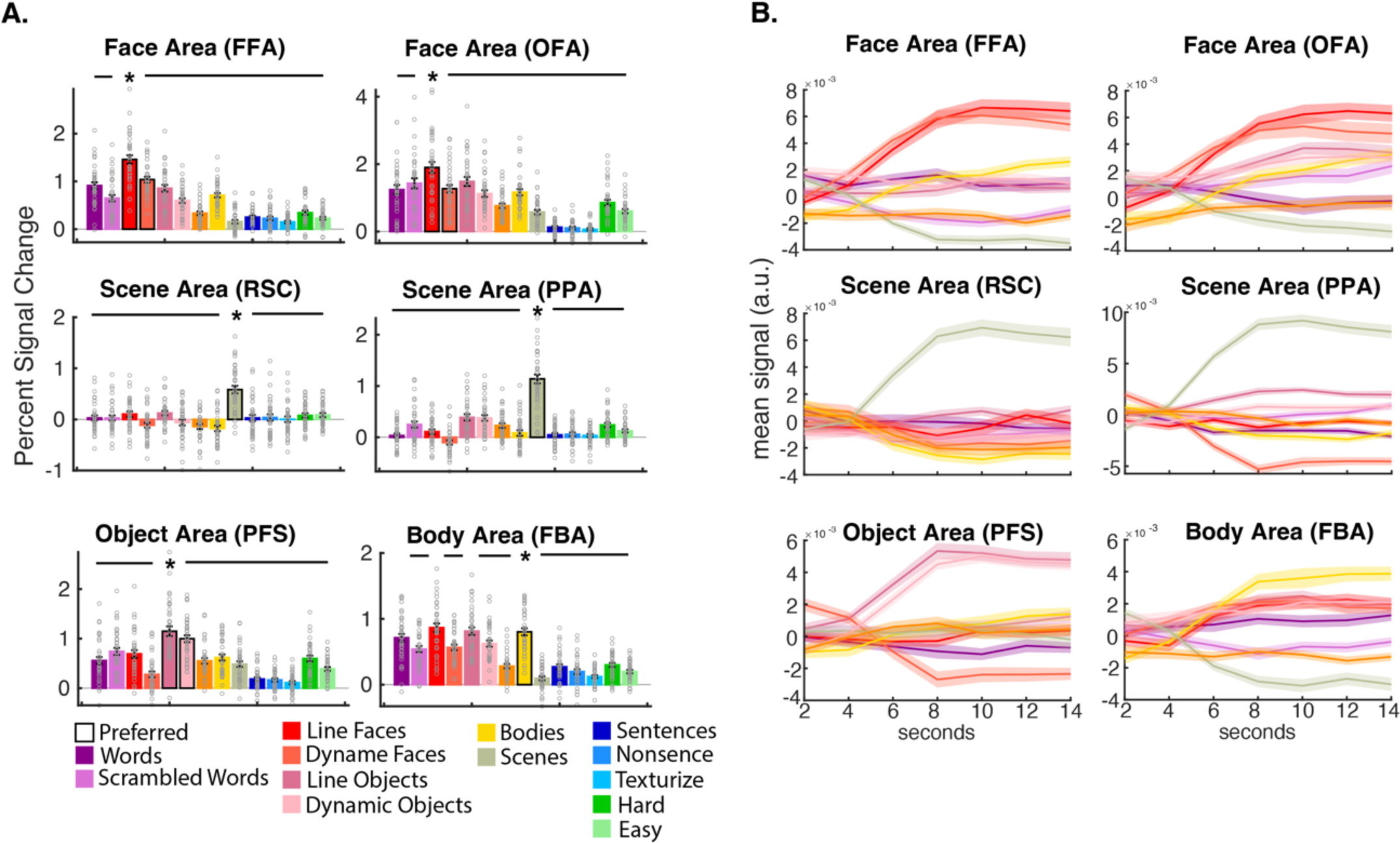
A. Functional profiles for left VTC high-level visual regions. The preferred condition(s) for each fROI is outlined in black. The mean percent signal change (with standard error bars) to various visual and non-visual conditions are plotted. Individual subject PSCs are shown with grey hollow circles. Significance is noted (*p<0.05, Bonferroni-holm) for the preferred category only, with a black lining showing all conditions significantly lower than the preferred category. B. Averaged time-course of each fROI’s responses to blocks of different experimental conditions. Dark line for mean across all subjects and shading for standard error. FFA: fusiform face area, OFA: occipital face area, RSC: retrosplenial cortex, PPA: parahippocampal place area, PFS: posterior fusiform sulcus, FBA: fusiform body area.

**Figure 3.**
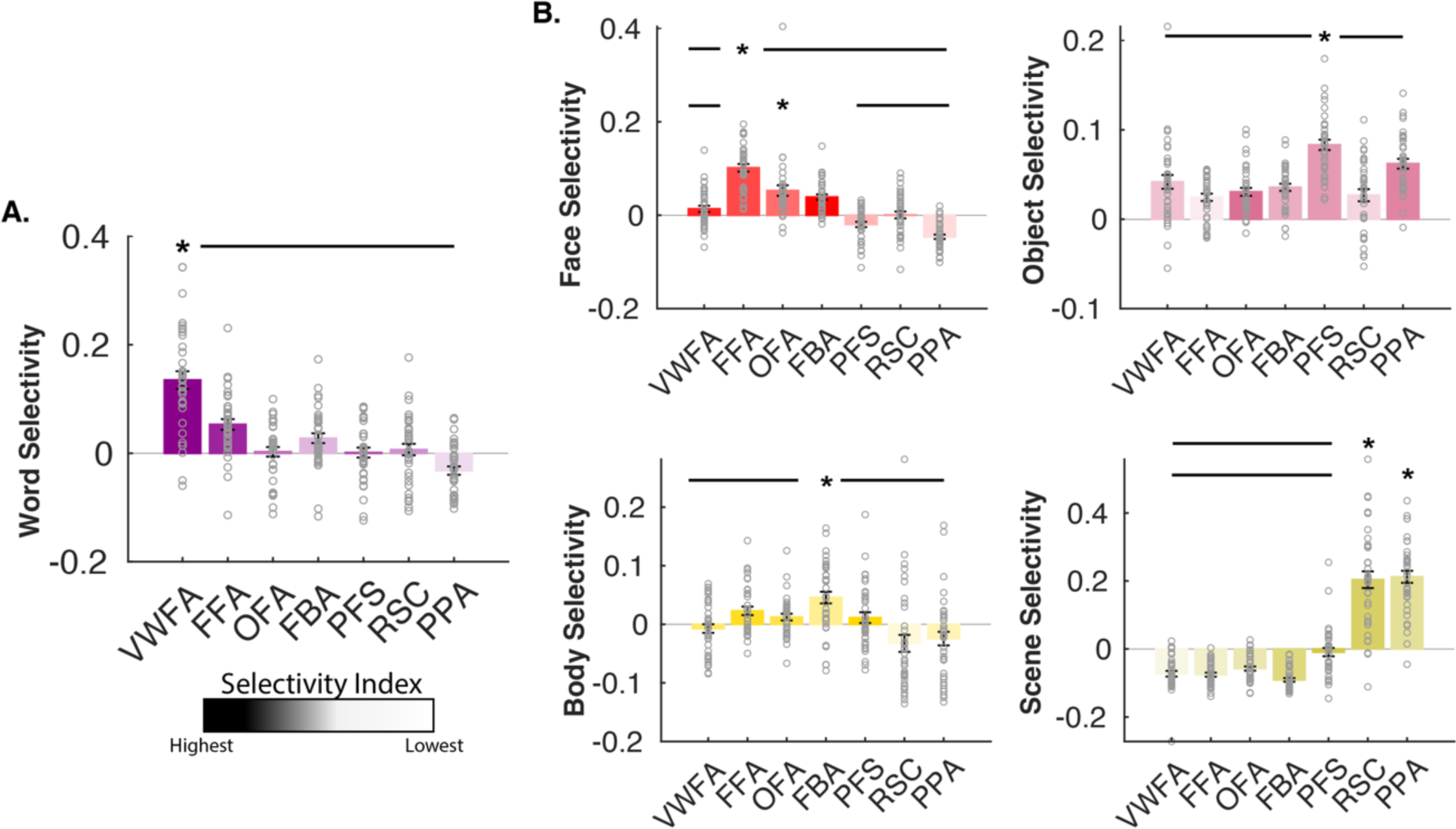
Category selectivity indices in the category-selective regions. For each visual category, we computed the selectivity indices (see **Methods**). Asterisks denote the specific category-selective regions associated with each category selectivity. Horizontal lines indicate the values are significantly (corrected) lower than that of the corresponding region.

Does the VWFA also show some preference for nonword stimuli, as shown in some prior work? Interestingly, when we examined the VWFA’s selectivity to non-preferred conditions, we found that in addition to the VWFA’s absolute preference for visual words, it also showed higher activity to objects (average of line-object and dynamic object vs. average others excluding the response to words: t(36)=3.17, p=0.003). Had we only used the dynamic localizer and examined the VWFA’s selectivity using meaningless scrambled objects as the control condition, commonly done in prior work, the VWFA would appear not only object, (t(35)=5.19, p=9.12x10^-6^), but also face (t(35)=1.99, p=0.054) and even body (t(35)=3.60, p=9.87x10^-4^) selective. However, these response patterns should not be taken as evidence against VWFA’s word selectivity. Instead, our results suggested that despite a distribution of responses to other visual categories, the VWFA shows an absolute highest word preference, highlighting the importance of comprehensively comparing the VWFA’s activation to a wide range of stimuli (**Figure 2 & 3; Table S1 & S3**).

One potential concern is that the observed results mainly reflected potential differences in scan parameters, motion, or SNR between experiments, thus we rescanned a subset of subjects on two additional runs of the static visual localizer. We replicate the main results even after matching these potential confounding factors (**Supplementary Results, Figure S4, and Table S5 & S6**).

### The functional landscape of the VTC

It is possible that the above results were biased by 1) the specific method (i.e., top 150 vertices) used to define fROIs; or 2) the predefined search spaces we used when defining the fROIs. Using the top 150 method results controls for the size of the fROIs in comparison, and also allowed us to identify a relatively small set of the most responsive voxels, which further avoided the overlap between regions to ensure spatial specificity. Indeed, when we explicitly quantified any overlap among the VWFA and other VTC fROIs when selecting the top 150 vertices before assigning them based on their most responsive category (see **Methods**), we found little overlap if any within the individual: an average of 13, 9, and 3 vertices overlapped between the VWFA and FFA, FBA, and PFS, respectively. Additionally, we found similar results (**Table S1**) when applying different criteria to identify the fROIs: selecting the top 10% of vertices within the search space and using a hard significance threshold (p<0.005) (see **Table S7** for descriptive information (e.g., number of subjects has the significant fROIs, size and the overlap between regions) for the fROIs defined with these other two methods).

Further, to complement the fROI analyses, which might miss selective responses outside the predefined search spaces and reveal little about the spatial spread of potential subregions of the word-selective areas, we examined voxel-wise selectivity to all visual stimuli, from posterior to anterior VTC in both fusiform and inferior temporal cortex (see **Methods**). Interestingly, we found selective responses for words, faces, and objects within the fusiform cortex, from the mid-fusiform and extending posteriorly (**Figure 4**). This suggested that selective voxels for different high-level conditions were close, but distinct, to each other within a relatively small swarth of the fusiform cortex. Moreover, only word-selective responses were found in the inferior temporal cortex, consistent with the notion that word selectivity is often found to be more lateral. Again, no reliable body-selective response was found even within the entire VTC; thus, the left FBA was excluded from further analyses.

**Figure 4.**
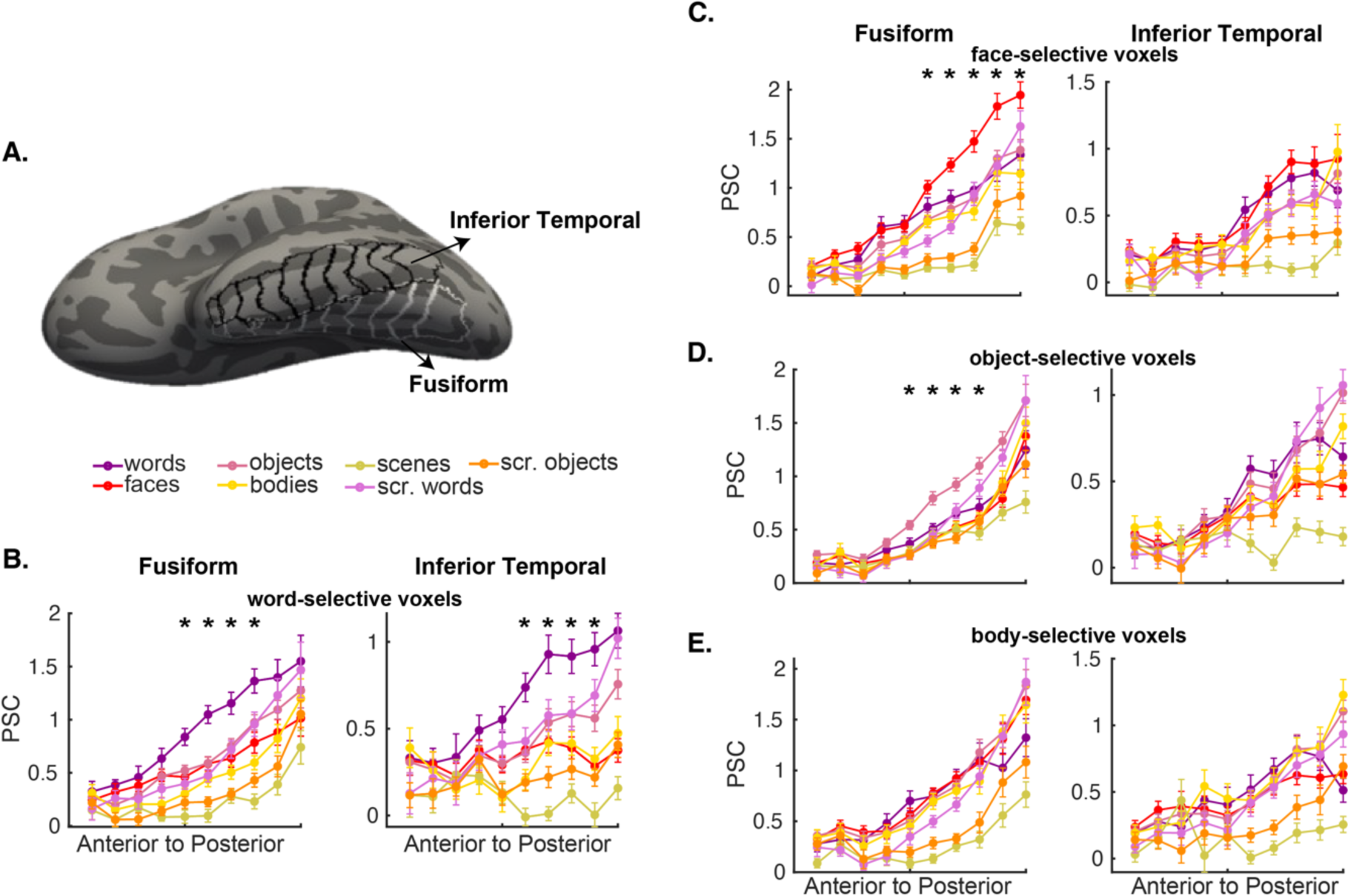
Categorical responses from the posterior to anterior left VTC. A. Interior temporal (black outline) and fusiform (white outline) parcel that comprise the VTC from the Desikan-Killiany parcellation. We divided each anatomical parcel into 10 equal sections from posterior to anterior. B-E. PSC to each of the visual conditions at each section along the posterior-to-anterior axis. The asterisk denotes that the PSC to the condition of interests is significantly higher than all the other conditions at a given location (corrected). Note that for faces and objects, we averaged PSC from the static and dynamic localizers.

### The VWFA, compared to adjacent VTC regions, selectively responds to auditory language, but is not a core part of amodal high-level language network

Next, we asked the extent to which the VWFA is multimodal, also responding to auditory language. We first investigated if the VWFA shows language-selective response by comparing its responses to English sentences (Sn) with nonword sequences (nonsense), presented auditorily (see **Methods**). Note that the nonword condition shares speech features like prosody and phonological processing with sentences, thus the difference indicates selective responses to high-level linguistic features (i.e., semantics and syntax). We found that only the VWFA showed significant language selective responses (Sn > Ns: t(35)=2.85, p=2.20x10^-2^), none of the other category-selective lVTC regions (FFA, PFS, OFA, RSC, PPA) differentiated between sentences and nonword sequences (all p > 0.05; **Table S4**). This language preference in the VWFA is also clear when examining the time-course of responses during the language task. (**Figure 5A**).

**Figure 5.**
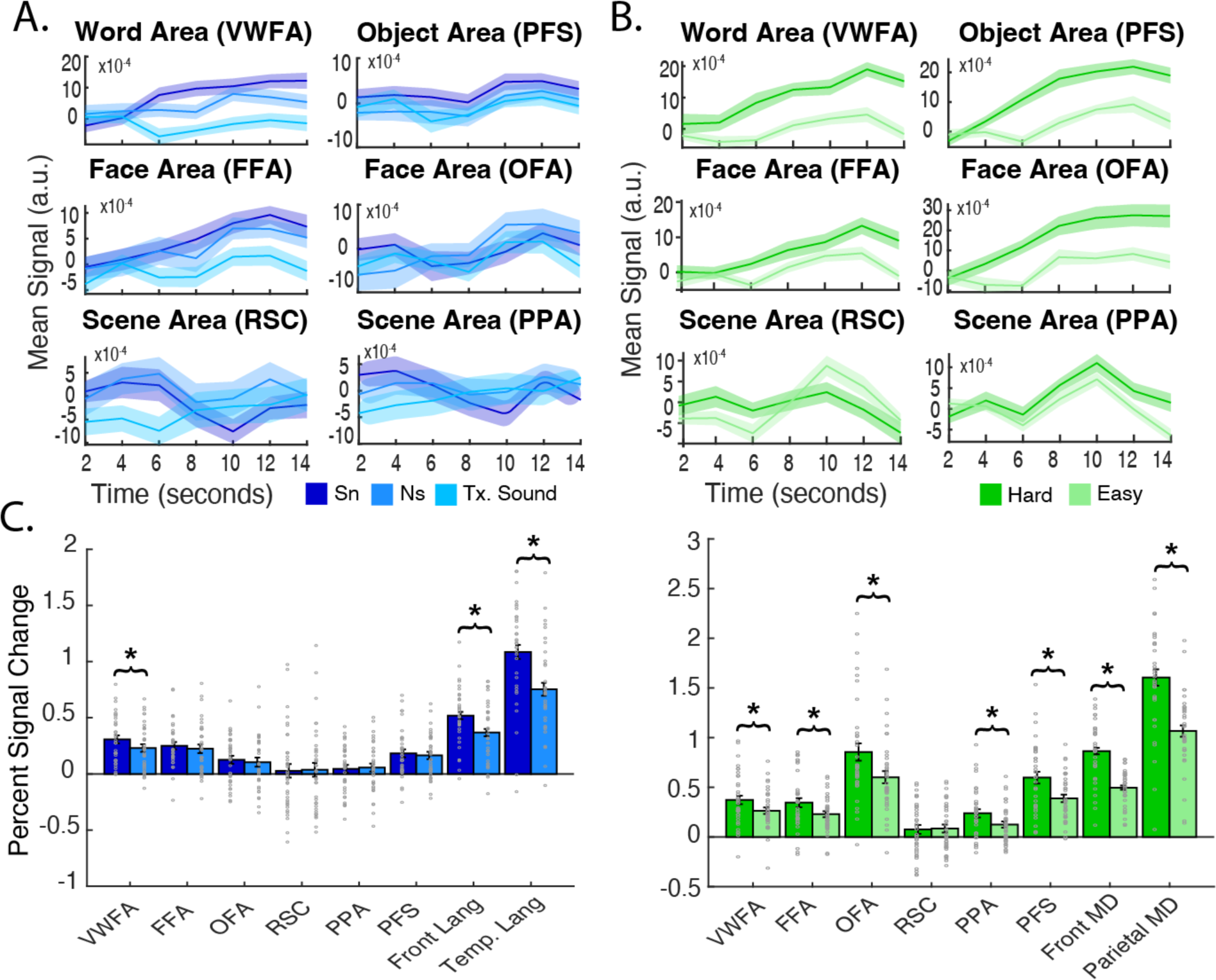
**A.** The time-course of each VTC fROI during the language task (averaged across all blocks). Dark line for mean across all subjects and shading for standard error. **B.** The time-course of each VTC fROI during the spatial working memory attentional load task (averaged across blocks). Dark line for mean across all subjects and shading for standard error. **C.** Mean percent signal change (with standard error bars) for each VTC fROI and the language fROIs to the language task (left) and the MD fROIs to the MD task (right). Individual subject PSCs are shown with grey hollow circles.

How does the VWFA respond to auditory language compared to the canonical language network (Fedorenko et al., 2010)? Unsurprisingly, the frontotemporal language fROIs (2 temporal and 3 frontal regions, see **Methods**) showed language selectivity (Sn > Ns, paired samples t-tests: collapsed across temporal fROIs: t(33)=7.43, p=1.54x10^-8^; and frontal fROIs: t(33)=4.59, p=6.21x10^-5^). Critically, as shown in **Figure 5C**, compared to canonical language fROIs, VWFA showed significantly lower activation to auditory language, regardless of conditions (Sn: temporal vs VWFA: t(33)=11.30, p=7.58x10^-13^; frontal vs VWFA: t(33)=5.21, p=9.90x10^-6^; Ns: temporal vs VWFA: t(33)=8.71, p=4.53x10^-10^; frontal vs VWFA: t(33)=4.51, p=7.80x10^-5^). If we had only explored language activation within the VTC, we may have concluded that the VWFA was in fact selective to amodal linguistic processing. However, as we have shown above, this language preference is dwarfed by responses within the canonical language network.

Additionally, not only does the VWFA respond lower to language than the language network, but it also responds significantly more to any visual category (even non-preferred ones) than auditorily presented linguistic stimuli (paired t-test between mean response to all non-preferred visual categories and auditory sentences: t(35)=2.83, p=0.0076), suggesting that the VWFA is primarily a visual region. Altogether, the higher activation to visual stimuli in general when comparing the visual vs. language responses of the VWFA further underscored its function as a high-level visual region specifically for processing orthographic stimuli.

As a comparison, we also examined the effect of domain-general attentional demands on the VTC, as frontoparietal multiple demand (MD) regions (see **Methods**) are in close vicinity of the language regions and previous studies also reported connectivity between the VWFA and dorsal parietal attention region. We confirmed that these MD regions demonstrated a significant attentional effect, measured by comparing the response to Hard vs. Easy conditions in a spatial working memory task: frontal MD: t(33)=10.24, p=9.02x10^-12^; parietal MD: t(33)=10.60, p=3.64x10^-12^. Critically, in contrast with the unique effect of language on the VWFA, almost all VTC fROIs (except for the RSC) were significantly modulated by attention (**Figure 5C**, right) and the attentional effect in the VWFA was similar to other fROIs (e.g., FFA) or at times even lower than other fROIs (e.g., PFS and OFA; see full pairwise comparisons in **Table S4**). Time course analyses of VTC fROIs during the MD task also confirmed this ubiquitous attentional effect (**Figure 5B**). Moreover, even though the spatial working memory task requires visual processing, we nevertheless found that the magnitude of the attentional load effect in the VWFA was significantly lower than the effect observed in frontal (MD vs. VWFA: Hard, t(33)=9.23, p=1.17x10^-10^; Easy, t(33)=5.73, p=2.17x10^-6^) or parietal MD regions (MD vs. VWFA: Hard, t(33)=13.47, p=5.80x10^-15^; Easy, t(33)=11.24, p=7.99x10^-13^). These results suggested that, in contrast to the linguistic effect, the modulation of attention is general within the VTC, highlighting the unique involvement of the VWFA during the auditory language processing.

### The VWFA is distinct from the basal temporal language area

As we have shown above, while VWFA uniquely showed some auditory language sensitivity, this response was lower than that in the core amodal language network. Lastly, we asked if there exist language clusters within the VTC that are distinct from the VWFA. We first examined the probabilistic map of the auditory language activation (**see Methods**). We found two clusters located in the left VTC that showed language selectivity (Sn>Tx): anterior language VTC (aLang-VTC, **Figure 6A**) and medial language VTC (mLang-VTC) (**Figure 6B**; similar clusters were observed for Sn vs. Ns, **Figure S5**; see also Figure **S6** for RH hotspots with similar attentional but weaker language activation). Using these two clusters as the search spaces, we defined subject-specific fROIs for mLang-VTC and aLang-VTC and examined their functional profile (see **Figure 6C)**. In the following section, we characterized the functionality of these two “language” regions quantitively at individual level in comparison to the VWFA and left amodal frontotemporal language network.

**Figure 6:**
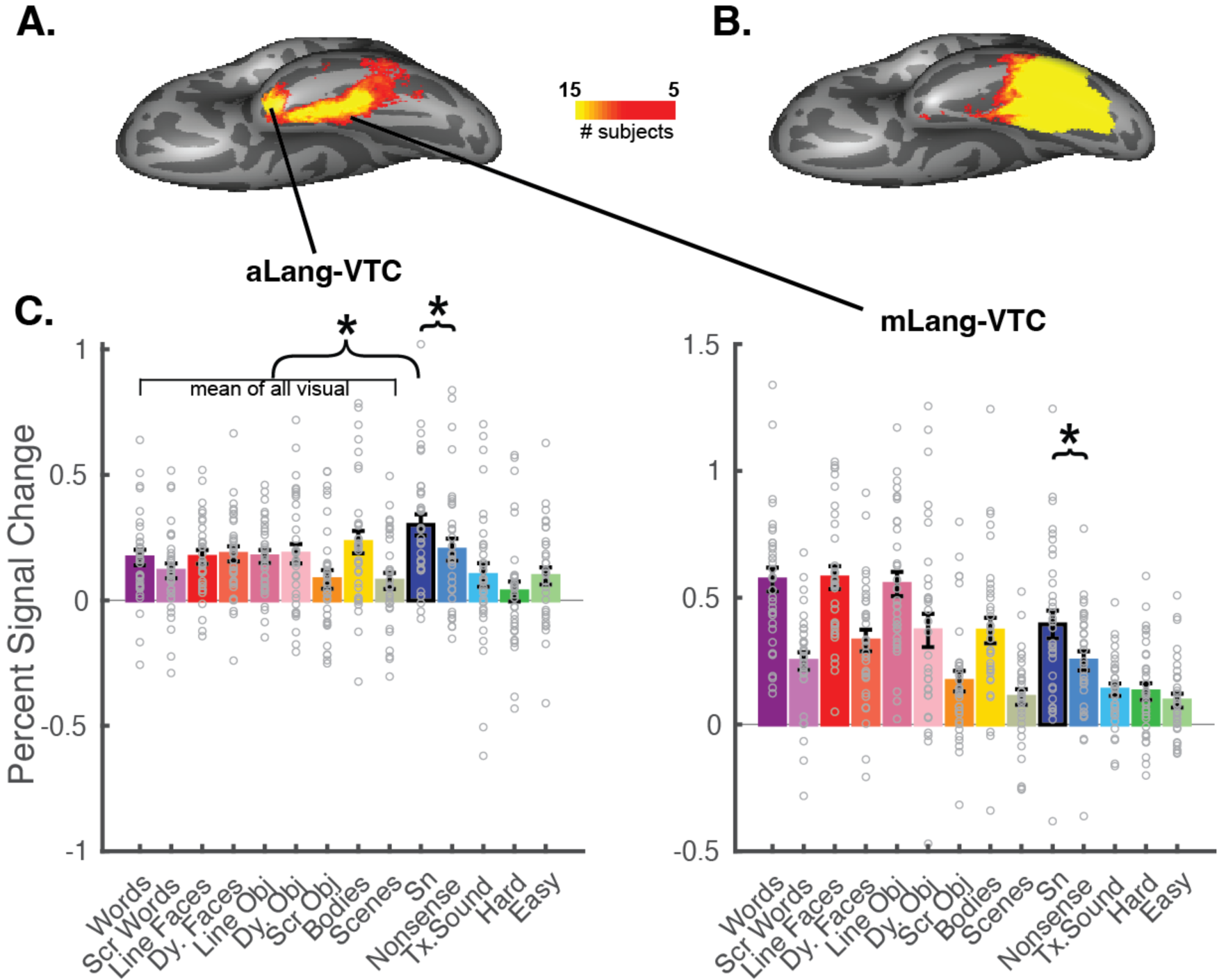
**A.** Probabilistic map for Sn>Tx showing subjects with overlapping activation during the language localizer within the VTC. For A and B, each subject’s statistical map was thresholded at p<0.01 and the resulting binarized maps were added together (minimum overlap = 5 subjects) (see **Methods** for details). **C.** Functional profile of the left aLang-VTC and mLang VTC fROIs. The mean percent signal change (with standard error bars) to various visual and non-visual conditions are plotted. The preferred category (i.e., auditory sentences) is outlined in black. Individual subject PSCs are shown with grey hollow circles. Brackets denote a significant difference between sentences and nonsense (paired t-test, *p<0.05). The dashed line denotes a significantly higher response to auditory sentences than visual categories on average for aLang (paired t-test, *p<0.05).

As expected, both the aLang-VTC and mLang-VTC were language selective (paired-samples t-tests (Sent>Ns): aLang-VTC, t(33)=3.84, p=5.32x10^-4^; mLang-VTC, t(32)=3.35, p=0.002). Critically, aLang-VTC’s response to sentences was significantly higher than its average response to all visual categories (paired-samples, two-tailed, t-tests: t(33)=3.45, p=0.0015), and it did not display a clear preference among these high-level visual categories (one-way rmANOVA of responses to the static localizer: F(2,64)=0.042, p=0.959; one-way rmANOVA of response to dynamic localizer: F(3,96)=7.024, p=2.55x10^-4^; post-hoc t-tests show significantly lower response to scenes, but bodies, faces, and objects are not distinguishable). In contrast, the mLang-VTC showed comparable responses to visual conditions as to auditory language (sentences vs. average response to visual stimuli: t(33)=0.45, p=0.66)). Importantly, however, just like aLang-VTC, mLang-VTC responded to different high-level visual stimuli equally and did not show a clear category preference (one-way rmANOVA of responses to the static localizer: F(2,66)=0.25, p=0.78; one-way rmANOVA of response to dynamic localizer: F(3,93)=11,67, p=1.47x10^-6^; post-hoc t-tests show significantly lower response to scenes, but bodies, faces, and objects are not distinguishable). These results, along with the observation that there is spatial overlap between the mLang cluster and the VWFA parcel, possibly explains why previous studies may have conflated the VWFA with these more anterior language regions. Here we show that there are two anterior language clusters that are functionally distinct from the VWFA (i.e. respond equally to all high-level visual categories) and may engage in abstract semantic processing generally.

## Discussion

Studies have continued to debate the existence and functional characteristics of the VWFA (see Dȩbska et al., 2023; Price & Devlin, 2003, 2011 for a review). Our study investigated the VWFA’s comprehensive response profile to a wide range of visual and non-visual stimuli and compared its neural signature to those of other spatially adjacent VTC regions. We found that while responding moderately to objects and faces, the VWFA’s response to visual words towers above its responses to all other high-level visual categories. Moreover, we found that while the VWFA is the only VTC region that showed sensitivity to auditory language, the VWFA is modality-dependent. The VWFA is primarily visual: its responses to non-word visual stimuli surpass its response to auditory language. In the following sections, we discuss defining a category-selective region (conceptually and methodologically), methodological discrepancies that might have contributed to the inconsistency regarding the VWFA’s function, the implications of non-word responses in the VWFA, and the hierarchical organization of the VTC: from posterior regions that respond to visual forms of the words (i.e., VWFA) to anterior areas associated with abstract semantics.

### The definition of the VWFA

Our results suggest that the muddled picture of the VWFA was at least partially driven by failure to take into account the individual variability of the VWFA as well as the intertwined nature of category-selective VTC regions (evident in the fROI map for each individual; **Figure S1**). Here we defined the VWFA with rigorous methodological considerations: the subject-specific approach (Fedorenko et al., 2010) to account for individual differences, different thresholds to select the candidate fROI voxels to examine the robustness of observed results, multiple control conditions to match either visual complexity or conceptual semantics for functional specificity, and simultaneously defining adjacent VTC regions to ensure high spatial specificity. While previous studies utilizing whole-brain group analysis observed no word-selective responses (e.g., Kherif et al., 2011; Vogel, Petersen, et al., 2012; see Price & Devlin, 2003, 2011 for a review), here we were able to localize a VWFA fROI in each individual that responded significantly higher to visual words than to all other visual and non-visual stimuli. Our results highlight the importance of defining the VWFA in each individual: when lumping all subjects together, either by implementing group analysis or drawing arbitrary spheres around predefined coordinates, we lose the precision and spatial resolution to identify word-selective voxels, as illustrated by the extensive overlap between the group-level probabilistic maps of different category-selective activations (**Figure S7**). This might be one reason why previous studies have reported that the VWFA responds to non-word stimuli like faces, objects, or symbols (Mei et al., 2010; Neudorf et al., 2022; Planton et al., 2017; Vogel, Petersen, et al., 2012).

Critically, we found that the VWFA was functionally different from other VTC fROIs: it showed minimal overlap with other VTC fROIs at the individual level, and also showed the highest responses to visual words versus other non-word conditions (e.g., nearly twice as much to words (average PSC 1.06 +/- 0.44) as to the second highest category (i.e., objects, average PSC of 0.58 +/- 0.35)). This aligns with the definition of a category-selective region that is domain-specific (Kanwisher, 2010). For example, the FFA shows higher activation to objects versus nonface conditions, but these responses are much lower than its responses to faces <u>(usually twice as low, Kanwisher et al., 1997; see Kanwisher (2010) for a discussion)</u>.

### Functional profile of the VWFA: activation for non-word stimuli

To what extent does the VWFA show preferential activity to other high-level visual categories? Answering this question may give us clues as to how this piece of cortex is able to process words. Previous work investigating the neuronal tunning of the ventral visual stream showed that the VTC is organized by underlying neuronal preferences for different visual and semantic features (e.g., fovea/peripheral bias, Hasson et al., 2002; simple geometrical features, Biederman, 1987; Sablé-Meyer et al., 2021; rectilinearity, Nasr et al., 2014; spatial frequency, Woodhead et al., 2011; spikiness, Bao et al., 2020; animacy and real-world size, Konkle & Caramazza, 2013). Therefore, some researchers proposed that the VWFA may emerge or be repurposed from part of another high-level visual region (see Dehaene & Cohen, 2007 for a review) that shares similar visual features with visual words. In this section, we discuss insights we gain from these non-word responses: that the cortical tissue later becomes word-selective also shows some sensitivity to local visual features like line segments and junctions, stimuli in the center visual field, and stimuli that encode abstract semantic information.

First, we found that while the VWFA responds more to words than other visual categories, its response to the scrambled words condition was surprisingly high (about as high to faces and higher than scenes and bodies). This result suggests that the VWFA’s preference for visual words may emerge from existing preferences for geometrical visual features such as line segments, junctions and contours (Lanthier et al., 2009; Szwed et al., 2009, 2011)

Second, the foveal hypothesis of the VTC proposes a medial-to-lateral dissociation in the ventral visual stream for processing peripheral and fovea stimuli, respectively. Consequentially, the lateral portion of the VTC houses both the FFA and VWFA, as visual words and faces are processed foveally (Hasson et al., 2002). We see that in our results as well, and we also find that the VWFA shows the least responses to scenes, fitting the lateral-to-medial functional division. We might then expect to see strong face responses in the VWFA (Plaut & Behrmann, 2011), as compared to other visual stimuli; however, we do not find that the VWFA responds more to faces than other high-level visual conditions in either static or dynamic localizer, except dynamic scrambled objects which only controls for low-level visual features such as color and edges, resembling the checkerboard condition of early VWFA functional localization (Cohen et al., 2002). This aligns with prior work showing overlap between word and face responses in the VWFA only when using fixation as a control condition to define VWFA (thus presumably including a large portion of lateral VTC rather than just the VWFA) Nestor et al. (2013).

Finally, among all non-word visual conditions, the VWFA responds highest to objects: it responds the second-highest to the line-drawing of objects and responses to objects (average of static and dynamic) are significantly higher than the average of all other non-word stimuli. This relatively high activation to objects was also observed in a previous study, where Ben-Shachar et al. (2007) reported that the VWFA responded second-highest to line-drawing objects, followed by false fonts. This may further explain difficulties of differentiating word-selective responses from objects, as noted in previous studies (e.g., Augustine et al., 2015; Kubota et al., 2019). Perhaps the representation of high-level visual objects is one of the other functions carried out by this piece of cortex. However, crucially, object responses within the VWFA were much lower than the VWFA’s responses to words and also much lower than the selectivity of face and object-preferring regions like the FFA and PFS, respectively, suggesting that while this piece of tissue may be able to represent objects, its main preference is for word stimuli.

### Word-selective responses along the posterior-to-anterior VTC

Studies have shown that distinct areas along the mid-fusiform and occipital temporal sulcus may be involved in processing different aspects of visual words (Cohen et al., 2002; Lerma-Usabiaga et al., 2018; Weiner et al., 2014; Zhan et al., 2023). For example, some work suggests a hierarchical organization of orthographic representation, becoming more abstract as one progresses more anteriorly (e.g., Vinckier et al., 2007). Specifically, the posterior region at the tail of the OTS, known as the pOTS (or VWFA-1; Grill-Spector & Weiner, 2014) is responsive to visual features of words (Lerma-Usabiaga et al., 2018) These groups have also identified a more anterior region at the mid-OTS (which also straddles fusiform gyrus and inferior temporal gyrus), known as the mOTS (or VWFA-2; Grill-Spector & Weiner, 2014) which is responsive to word form and language units (Lerma-Usabiaga et al., 2018). Other studies (see Jobard et al., 2003 for a review) also report more anterior word-selective regions but it remains unclear whether these regions represent orthography/script or whether they represent general visual semantics/abstract concepts (e.g. another object area) because previously used fMRI contrasts were limited to words versus meaningless letter-like stimuli.

In the current study, the VWFA parcel that serves as our search space covers the mOTS and extends anteriorly. When we examined the full gradient of responses along the VTC and compared words versus both scrambled words and objects (to control for abstract concepts), we found that word-selective voxels extended from mOTS to a more anterior mid-fusiform region that is specialized for orthographic lexicon (Wimmer & Ludersdorfer, 2018). In line with the idea of this orthographic selectivity that relies on the recognition of letter sequences of recurring word parts (rather than the meaning of words), previous studies have shown that the VWFA is sensitive to orthographic regularity (e.g., frequency of letter bigrams or trigrams) rather than lexical status (distinguishing pseudowords from real words) (Binder et al., 2006; Vinckier et al., 2007). General lexical or visual semantics, interestingly, might be associated with an even more anterior cluster (Lerma-Usabiaga et al., 2018), which we will discuss further below.

### Amodal linguistic activation in the left VTC

Another goal of the current study was to determine to what extent the VWFA responds to auditory language. We observed higher activation of auditory sentences vs. nonsense speech within the VWFA; but perhaps more importantly, the high-level linguistic response was significantly lower than the VWFA’s response to written language (i.e., visual words) and even to non-preferred visual categories. Further, responses to auditory language within the VWFA were dwarfed by the language responses of the frontotemporal language regions. Altogether, our results suggested that the VWFA is dominated by visual stimuli, rather than a modality-independent language-related region as claimed in a recent review (Dȩbska et al., 2023); or at least, our result suggests that VWFA might function differently than canonical language regions as it is in fact more tuned for visual aspects of language.

Interestingly, while not part of the core language network, the VWFA is the only a-priori-defined VTC region that shows high-level linguistic sensitivity. This tuning is likely due to coactivation between the VWFA and frontotemporal language regions via privileged connectivity between them (i.e., Connectivity Hypothesis, Mahon & Caramazza, 2011; with empirical evidence provided by Saygin et al., 2016; Li et al., 2020; Stevens et al., 2017). Similarly, Buckner et al. (2000) observed a repetition priming effect for auditory words on the inferior temporal cortex and they further proposed that the top-down effect from frontal regions might account for this auditory activation, likely via connectivity between the VWFA and frontal regions (e.g., Bouhali et al., 2014; Saygin et al., 2016; Thiebaut de Schotten et al., 2014). Therefore, the connectivity between the VWFA and language regions may prepare that piece of the cortex for language-related stimuli, and with the visual experience of written language (i.e., orthographic stimuli), it further tunes for and becomes functionally selective for visual words as shown in our results here. This aligns with the idea that both connectivity and experience further shape and constrain its functional specialization (Bedny, 2017). Conversely, when no visual input is available, the VWFA functions as a language region that demonstrates sensitivity to e.g. grammatical complexity (Kim et al., 2017).

In addition to privileged connectivity with the language network, the VWFA also connects with the frontoparietal attentional regions (Chen et al., 2019; Vin et al., 2024; Vogel, Miezin, et al., 2012)). This provides one possible explanation for the VWFA’s activation in e.g. non-orthographic tasks (Mano et al., 2013; Starrfelt & Gerlach, 2007), which might be due to top-down feedback through VWFA’s connectivity (Planton et al., 2019), either by explicit task manipulations and demands (Moore & Price, 1999; Phillips et al., 2002; Wang et al., 2018; White et al., 2023) or long presentation times (1.5s, Vogel, Petersen, et al., 2012). However, in the current study, by implementing a 1-back task with fast presentation of visual stimuli (500ms), we demonstrated the VWFA’s dominant role in rapid and efficient visual word perception (Grill-Spector et al., 2000; Thorpe et al., 1996). And further, we show that most VTC regions are engaged more for hard versus easy spatial working memory, suggesting that the VWFA is not unique in this regard. Taken together, we suggest that the VWFA’s responses to auditory language and cognitive effort are the result of top-down influence and connectivity from other cortices, rather than robust neural preferences to these stimuli.

### A language cluster in VTC that is distinct from VWFA

Interestingly, our exploratory analysis showed that anterior to the VWFA (and anterior to VWFA-1/p-OTS and VWFA-2/mOTS), there are two language clusters within the left VTC that show linguistic selectivity, i.e. higher responses to auditory English sentences than to nonsense speech. Importantly, however, by directly comparing the functional response profile of these two regions to the VWFA as well as other VTC regions, we found these two language regions do not distinguish between different visual categories (including words, unlike the VWFA). In fact, the anterior cluster (aLang-VTC) is seated at the tip of the inferior temporal and fusiform gyrus, likely corresponds to the temporal pole region that was previously associated with language comprehension and semantic processing (Binder et al., 2009; Purcell et al., 2014) regardless of modality. Consistent with this, our results showed that aLang-VTC prefers auditory sentences more than visual stimuli. This was the only region within VTC that showed higher preferences for auditory sentences than other stimuli.

The more medial and posterior region, the mLang-VTC, however, showed comparable language activation to visual activation. This cluster might be the “basal temporal language area (BTLA)” that is situated between the left temporal pole and the VWFA according to Purcell et al. (2014). Note though, the role and even the anatomical location of the BTLA remains unclear and the term “basal temporal language area” is often used to refer to any or all language areas in the basal temporal lobe. Nevertheless, our results provide some insight into the role of this region: instead of specifically serving as the interface between semantics and orthography per se (Purcell et al., 2014), this multimodal region may play a role in the semantic processing of both words and other visual categories (Jobard et al., 2003; Thompson-Schill et al., 1999). This notion is supported by observations that the resection of the BTLA shows the strongest association with deficits in object naming compared to other VTC sites (Wilson et al., 2015).

### Limitations and Future directions

The present study systematically examined the role of the precisely defined VWFA and provided a clear characterization of the nature of its orthographic selectivity by looking at its activity in response to visual words, other non-word stimuli, and spoken language. However, limitations and open questions remain. First, as an effort to estimate a more comprehensive response profile of the VWFA, we scanned participants with multiple tasks. Ideally, we would want to test the function of the VWFA in a single task that includes as many conditions as possible to better compare between conditions. While we matched the scanning parameters (and most of the tasks were scanned within the same session), future studies should test the functionality of the VWFA with rich stimuli in the same task setting to further verify our results. Second, we complemented our fROI analysis with a gradient analysis to further probe the anatomical location of the word-selective voxels. However, it remains unclear whether the voxels we found in the mid-fusiform and inferior temporal cortex belong to one single cluster or if they are two distinct/separate clusters. Third, while we performed a gradient analysis of category-selectivity along the VTC, our experiment was not set up to explore the progression of abstract word-form representations along the VTC. And so it remains unclear whether the VWFA processes words vs. consonant strings/pseudowords differently (Dehaene et al., 2005; Vinckier et al., 2007) (although see Baker et al. (2007)for a comparably defined VWFA responding similarly to words and meaningless consonant strings) or whether the VWFA differentiates stimuli with different levels of orthographic regularities. Fourth, does the VWFA respond differently to visual words compared to other human-invented visual signs (e.g., numbers, traffic signs)? For example, Changizi et al. (2006) noted that there are common line configurations in human-invented visual signs which are not only presented in orthography, but also in other visual symbols, therefore the possible sensitivity of the VWFA or other VTC regions should be explored with respect to these symbols. Finally, what does the VWFA do prior to literacy, and what computations is this neural tissue capable of? Our hope is that this paper will clarify the role of the VWFA as a high-level visual, category-specific region for words, allowing research to move forward with understanding how this region develops its specialization as a fascinating example of uniquely human neural cognition.

## Methods

### Participants

Sixty-three healthy adults were recruited from The Ohio State University (OSU) and the local community. As part of an ongoing project exploring the relationship between functional organization of the human brain and connectivity, participants completed a battery of functional tasks in the scanner. In the current study, all 63 individuals completed at least one run of at least one of the functional tasks of interest: a VWFA localizer (N=56), a dynamic visual localizer (N=52), an auditory language task (N=48) and a spatial working memory task (N=57). The full sample of a given task was used to generate probabilistic maps for a given functional contrast. Critically, only individuals who completed two runs of the VWFA localizer and the dynamic localizer as well as at least one run of the other two tasks (N=37, 22 female, mean age: 24.57 age range: 18.01-55.66, 4 left-handed) were included in the main analysis, to ensure independence when defining the VTC category-selective regions and investigating their functional responses. When exploring the activation within the language and MD network, 3 individuals with only one run of the language task were further excluded. All participants had either normal vision or corrected-to-normal vision and reported no neurological, neuropsychological, or developmental conditions. The study was approved by the Institutional Review Board at OSU and all participants provided written consent.

### fMRI tasks

#### VWFA localizer (visual)

We used a VWFA localizer task (Saygin et al., 2016) to define the VWFA. Briefly, static images of words, scrambled words, objects, and faces were presented in blocks. In our critical visual word condition, a single printed word in black text color was shown and all words were nouns. Each word was presented on a black grid inside a white rectangular box, and for the scrambled words condition, each block of this grid was randomly arranged, to preserve low-level visual features of words (lines, curves intersections). Line drawings of faces and objects (also presented on a grid) were implemented to control for the effects of visual similarity, complexity, and abstract semantic meaning. All stimuli were displayed on a white rectangular box and a gray grid was superimposed on the stimuli to match edges that are present in the scrambled words condition (all stimuli are available for download at https://www.zeynepsaygin.com/ZlabResources.html). For this localizer, participants were asked to perform a one-back task. In each block, 26 stimuli (including 2 repetitions) of the same category were presented one-by-one for 500ms followed with a 193ms ISI. Each run contains 4 experimental blocks for each condition and 3 fixation blocks and 2 runs of the VWFA localizer were collected for each participant.

#### Dynamic localizer (visual)

To define other VTC category-selective regions that are adjacent to the VWFA, we used a dynamic visual localizer task where participants were asked to passively view video clips (with natural colors) from faces, objects, bodies, natural scenes and scrambled objects (Pitcher et al., 2011). Briefly, the face condition included faces of young children dancing and playing, the object condition included different moving objects (e.g., round block toys swinging), and the body condition showed different body parts (e.g., legs, hands; no faces) naturally moving. These clips were filmed on a black background. Scrambled objects were generated by scrambling each frame of the object movie clip into a 15-by-15 grid. Scene conditions were recorded from a car window while driving through a suburb. Six 3-s video clips from the same category were shown in each block (i.e., 18s per block) and 2 blocks of each experimental condition were presented in each run (alternating with a palindromic manner) with 3 rest blocks with full-screen colors alternating at the beginning, middle and the end of each run. The order across participants was randomized.

#### Language task (auditory)

A language task (Fedorenko et al., 2010) was used to investigate how the VWFA, as well as other VTC regions respond to the lexical and structural properties of auditory language. Participants listened to blocks of meaningful sentences (Sn), nonsense sentences (Ns, controlling for prosody but constructed from phonemically intact nonsense words), and texturized (degraded) speech (Tx, controlling for low-level auditory features). Each run consisted of 4 blocks of each condition and three 14-second fixation blocks. Each block contained three trials (6 seconds each); each trial ended with a visual queue to press a button. Language selective response is usually characterized by Sn>Tx or Sn>Ns, the former targets both linguistic and speech-related processing and the latter specifically targets high-level lexico-semantic information. Note that, we also defined the frontotemporal language regions (along with frontoparietal attentional regions, see below) so that we can compare the response of the VWFA to auditory languages with that of the canonical language-selective regions.

#### Spatial working memory task

As a comparison of the high-level linguistic effect, a spatial working memory task (Fedorenko et al., 2013) with blocked easy and hard conditions was used to examine the effect of attentional load on the VTC regions. The hard condition elicits higher activation than the easy condition due to the attentional load, which is the neural signature for the domain-general multiple-demand (MD) network(Blank et al., 2014; Fedorenko et al., 2013). In each trial, participants viewed a grid of six boxes. For the easy condition, three blocks would flash blue sequentially, and participants were expected to remember which of the boxes in the grid had been colored. Two grids would then appear, one with the same pattern of blocks colored as the previous sequence (the match) and the other with a different pattern of blocks highlighted (no-match), and participants used a button press to indicate the match. For the hard condition, 3 pairs of 2 blocks would flash sequentially, increasing the amount of information the participant needed to attend to and remember. Each run consisted of 3 blocks of 2 trials per condition, separated by 16-second rest blocks with a fixation on the screen. The order of conditions within each block was randomized across participants.

### Data acquisition and Preprocessing

All structural and functional MRI images were acquired on a Siemens Prisma 3T scanner (at the Center for Cognitive and Behavioral Brain Imaging (CCBBI) at the OSU) with a 32-channel phase array receiver head coil. All participants completed a whole-head, high resolution T1-weighted magnetization-prepared rapid acquisition with gradient echo (MPRAGE) scan (repetition time (TR) = 2300 ms, echo time (TE) = 2.9ms, voxel resolution = 1.00 mm^3^). A semi-automated processing stream (recon-all from FreeSurfer) was used for structural MRI data processing. Major preprocessing steps include intensity correction, skull strip, surface co-registration, spatial smoothing, white matter and subcortical segmentation, and cortical parcellation. For functional tasks, the VWFA localizer task was acquired with echo-planar imaging (EPI) sequence, TR=2000ms, TE=30ms, and 172 TRs. For N=50 participants, 2-mm isotropic voxels were acquired with 100 × 100 base resolution; 25 slices approximately parallel to the base of the temporal lobe to cover the entire ventral temporal cortex. For the rest of the participants, slightly larger voxels (2.2-mm isotropic) were acquired for whole brain coverage (54 slices) with the same number of TRs to cover the whole brain. All other tasks were also collected with EPI sequence, but with TR=1000ms, TE=28ms, voxel resolution of 2x2x3 mm^3^, 120 × 120 base resolution, 56 slices for the whole-brain coverage, and 244 TRs for the language localizer, 234 TRs for the dynamic localizer, and 400 TR for the spatial working memory localizer. To ensure that our results are not due to differences in scanning acquisition parameters, we collected an additional 2 runs of the VWFA task on 8 participants in our sample using identical parameters to the other 3 tasks, finding similar results (see **Supplementary Results 3**) for all analyses.

All tasks were preprocessed in the same manner. Functional data were motion corrected (all timepoints aligned to the first timepoint in the scan) and timepoints with greater than 1mm total vector motion between consecutive timepoints were identified (and later included in first level GLM as nuisance regressors). Data were distortion corrected, detrended, spatially smoothed (3mm FWHM kernel for the static visual and 4mm for the other tasks and 4mm for the static visual task with identical scanning parameters), and then registered from functional space to anatomical space (using bbregister from FreeSurfer). A block design with a standard boxcar function (events on/ off) was used to convolve the canonical HRF (standard g function, d = 2.25 and t = 1.25), and experimental conditions for each task were included as explanatory variables. Six orthogonalized motion measures from the preprocessing stage were included as additional nuisance regressors for each task individually. Resulting beta estimates, contrast maps and preprocessed resting-state data were registered from functional, volumetric space to anatomical, surface space (using FreeSurfer’s mri_vol2surf with trilinear interpolation), and then to FsAverage surface space (using FreeSurfer’s mri_surf2surf) for subsequent analyses.

### Definition of the functional region of interest (fROI)

Given the individual variability in the precise location of high-level visual regions across individuals, we used the group-constrained subject-specific method (Fedorenko et al., 2010) to define subject-specific fROIs. Previously defined atlas (parcels) that show the typical location of the regions across large samples of adults were used as our search spaces: for VTC parcels, VWFA was from Saygin et al. (2016) and all other parcels were from Julian et al. (2012) (except for lOFA and lFBA, see below); language functional parcels were from Fedorenko et al. (2010) (we used an updated version based on 220 participants) and MD functional parcels were original parcels from Fedorenko et al. (2013) (we used an updated version based on 197 participants) (see https://tinyurl.com/5e4tp67w for more details for language and MD parcels). For language and MD fROIs, we averaged results by lobes and reported results in frontal and temporal language regions, as well as frontal and parietal MD regions. All parcels were originally in volumetric spaces and moved to FsAverage surface with the same method mentioned above. For each subject’s fROI, we selected the most significant/responsive voxels within a given search space using the statistical maps of the contrast of interest. **Table 2** shows the corresponding localizer tasks and contrasts we used to define fROIs. Importantly, the most significant 150 voxels were selected and any voxels that responded significantly to multiple conditions were assigned to the contrast that they were most responsive to. To confirm our results were robust regardless of ways to define the fROIs, we also used two other widely used methods to select the voxels: a. applying a hard threshold to select the significant voxels; b. choosing the top 10% voxels within the search space. We replicated our main results in **Supplementary Results 1**. After fROIs were defined with one run of data, an independent run of data was used to extract the percent signal change (PSC) within each individual’s fROIs to all conditions of interest (**Table 2**). Any PSC values that exceed +/- 3 standard deviation across subjects were marked as outliers and removed from subsequent analyses.

**Table 2.**
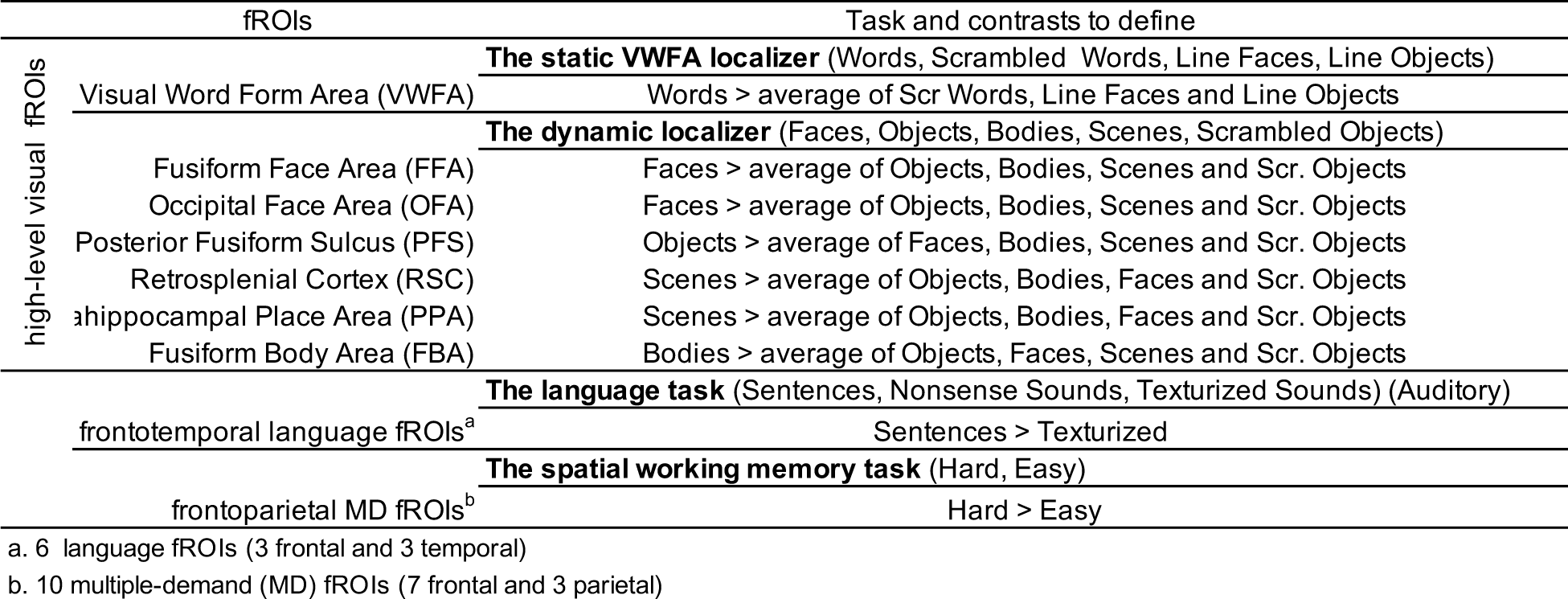
Tasks and contrasts to define the functional regions of interest.

Based on the PSC values, we additionally calculated category selectivity indices to test for their preferred and any unexpected non-preferred. Specifically, for their preferred category selectivity, the difference between the response to the preferred condition and the average of all other conditions was calculated; for the selectivity of the non-preferred category, we calculated the difference between the condition of interest vs. all other conditions after excluding the response to their preferred category. For example, the VWFA was tested for a preference for objects (average of dynamic and static objects – average of faces (dynamics and static), bodies, scenes, scrambled words, scrambled objects divided by the sum of all conditions)

In addition to canonical frontotemporal language-selective regions, to further explore the functional properties of the clusters within the VTC that respond to auditory language and compare them to the VWFA, we additionally defined two VTC language fROIs (mLang-VTC and aLang-VTC). Specifically, we first created the search spaces for the fROIs based on the Sn-Tx probabilistic map (see below). The same GcSS method was then used to define subject-specific mLang-VTC and aLang-VTC fROIs by selecting the top 150 language-responsive (Sn>Tx) voxels within the medial and anterior parcels we created below. Note that we identified these two fROIs along with the other category-selective VTC regions so that we could assign vertices to the condition to which they showed the strongest response (to achieve better spatial specificity, e.g. the mLang-VTC parcel overlapped with the VWFA parcel but the fROIs selected within these parcels showed the highest responses to the category of interest and do not overlap).

### Probabilistic map

As a sanity check and to justify the use of the functional parcels from independent studies as our search spaces when defining the functional regions of interest (see **Definition of the functional region of interest** below), we created probabilistic maps for different functions of interest. Using the contrast maps from the GLM analysis for each subject, we created probabilistic activation maps for different contrasts of interest. Specifically, for each subject, the statistical map based on all available runs of a task from a given contrast was minimally thresholded at p<0.01 (uncorrected) and resulting maps were binarized. The binarized maps of all subjects were then summed. This resulted in a map where the value at each vertex indicates the number of subjects who showed significant activation for that contrast at that location. We presented any vertex that showed a consistent significant effect of the tested contrast across at least 5 subjects (which is approximately 10% of the participants - the exact percentile varied because the total number of subjects who completed each task was slightly different).

The probabilistic map for words (words vs. the average of other conditions in the task) was made with the static VWFA localizer. The probabilistic maps for other high-level visual categories were created with the dynamic localizer: face (faces vs. objects), object (objects vs. scrambled objects), body (bodies vs. objects), and scene (scenes vs. objects) (**Figure S7**). Our probabilistic heat maps agreed well with most of the reference parcels with a few exceptions: lFFA parcel from Julian et al. (2012) was too small to cover the hot spots of face-selective activation around the left fusiform gyrus that we observed in our face probabilistic map (**Figure S7**). Therefore, we flipped the rFFA parcel to the left hemisphere. Additionally, given the original lOFA and lFBA parcels were missing the hot spots in the face and body probabilistic maps, we further created new lOFA and lFBA parcels based on the probabilistic maps. Specifically, we chose vertices that were consistently significant (p<0.01 at the individual level) across at least 5 subjects; and we used the cluster function from FSL (5.0.10) to obtain the cluster pass this criterion. Our main results use the updated lFFA, lOFA and lFBA parcels.

Moreover, to explore possible amodal language and attentional load activation within the VTC, we used the auditory language and spatial working memory tasks to generate probabilistic maps within the VTC for language (contrasts: Sn>Tx) and attentional load (contrast: hard vs. easy). While Sn<NS speech yielded similar hotspots as the Sn>Tx (**Figure S5**), we used this relatively liberal contrast so that the resulting parcel serves as a loose spatial estimation that allows us to search any possible language-selective voxels (see below). Just as how we made the lOFA and lFBA parcels, the summed map across individuals was thresholded so that at least 5 subjects showed significant Sn>Tx at a given vertex, which resulted in two continuous parcels (with the cluster function from the FSL) at the medial lVTC (mLang-VTC, anterior to the fusiform gyrus) and superior anterior lVTC (aLang-VTC).

### Gradient analysis

We complemented the fROI analysis with a gradient analysis, where we explored the category selectivity of the entire VTC. Specifically, using the Desikan-Killiany cortical parcellation(Desikan et al., 2006), we first divided the VTC into the fusiform and the inferior temporal cortex, and then within each anatomical parcel, we created 10 equal sections from the posterior to the anterior (i.e., the size of each slice along the y-axis is equal to the one-tenth of the parcel along that axis). With one run of the data, we identified the potential category-selective voxels within each section with a hard threshold (p<0.01), and in the independent run, we extracted those voxels’ responses (i.e., PSC) to each of the high-level visual categories in each section (for faces and objects, responses in the static and dynamic localizers were averaged). Note that we did not include scene selectivity in this analysis as this scene-selective in the VTC is usually more medial to all of the other VTC regions (i.e., medial to the fusiform cortex).

### Statistical analysis

Mostly, we used paired t-tests (two-sided) to compare between conditions or fROIs to establish response selectivity and specificity. Bonferroni-Holm multiple comparisons was used to correct for the number of comparisons for each analysis. Moreover, one-way repeated-measure ANOVA (rmANOVA) were used to compare category selectivity among VTC fROIs as well as to examine the visual category specificity of the aLang-VTC and mLang-VTC. Follow-up post-hoc paired t-tests were used to identify the differences (and corrected). Paired t-test was run with the Matlab ttest function and all ANOVAs were completed in RStudio (V 1.4.1717) using the anova_test function, with the subject number and fROI (or condition where applicable) as within-subject factors.

## Data and code availability

The data and code used in the current study are available from the corresponding author upon request.

## Author Contributions

J.L.: conceptualization, formal analysis, methodology, writing-original full draft & editing; K.H.: conceptualization, formal analysis, writing-original draft (sections) & editing; Z.M.S conceptualization, supervision, writing-review & editing.

## Acknowledgments

We appreciated the participation of our subjects. We would like to thank the Saygin Developmental Cognitive Neuroscience Lab members for helping with data collection and providing suggestions and feedback. We would like to acknowledge the support from the Center for Cognitive and Behavioral Brain Imaging (CCBBI) and the Ohio Supercomputer Center (OSC). This research is supported by the Alfred P. Sloan Fellowship (to Z.M.S.) and NSF Graduate Research Fellowship Program (DGE-1343012) to K.H..

## Declaration of generative AI and AI-assisted technologies in the writing process

During the preparation of this work, the author(s) used ChatGPT and Grammarly in order to check grammar. After using these tools/services, the author(s) reviewed and edited the content as needed and take(s) full responsibility for the content of the publication.

