## Supplementary Information for "Demystifying the Visual Word Form Area: Visual and Nonvisual Response Properties of Ventral Temporal Cortex with precision fMRI"

**Figure S1.** Subject-specific fROIs for the left and right ventral temporal cortex (VTC)
**Figure S2.** rVWFA functional response profile
**Figure S3.** Functional response profiles for right VTC category-selective fROIs.
**Table S1.** Comparing each fROI's preferred condition vs. other conditions for all methods of defining fROIs (LH).
**Table S2.** Comparing each fROI's preferred condition vs. other conditions for all methods of defining fROIs (RH).
**Table S3.** Comparing category selectivity among left VTC
**Table S4.** Language-selective effect and attentional effect in the left VTC fROIs
**Supplementary Results.** Replication for new VWFA parameters.
**Table S5.** Comparing each fROIs PSC to its preferred condition vs. other conditions (matching parameter subset)
**Table S6.** Language- and attentional-selective responses in the left VTC fROIs (matching parameter subset)
**Figure S4.** Functional profile for left and right VTC fROIs in 8 subjects who completed the static visual localizer with scan parameters matching all other functional localizers
**Figure S5**. Probabilistic maps for the language-selective response, defined with Sentences (Sn) > Nonsense Sounds (Ns), within the bilateral VTC
**Figure S6.** Probabilistic maps for language and load-based attentional responses in the right VTC.
**Figure S7.** Probabilistic maps for faces, objects, scenes, and bodies, created with the dynamic localizer
**Table S7.** Descriptive information for fROIs when defined with a hard threshold and top 10%

**Figure S1: Subject-specific VTC fROIs (VWFA, FFA, OFA, PFS, PPA, RSC, FBA) for all subjects in the left (top) and right hemisphere (bottom).** fROIs were created using the top 150 most significant voxels within parcel search spaces (see main text methods).

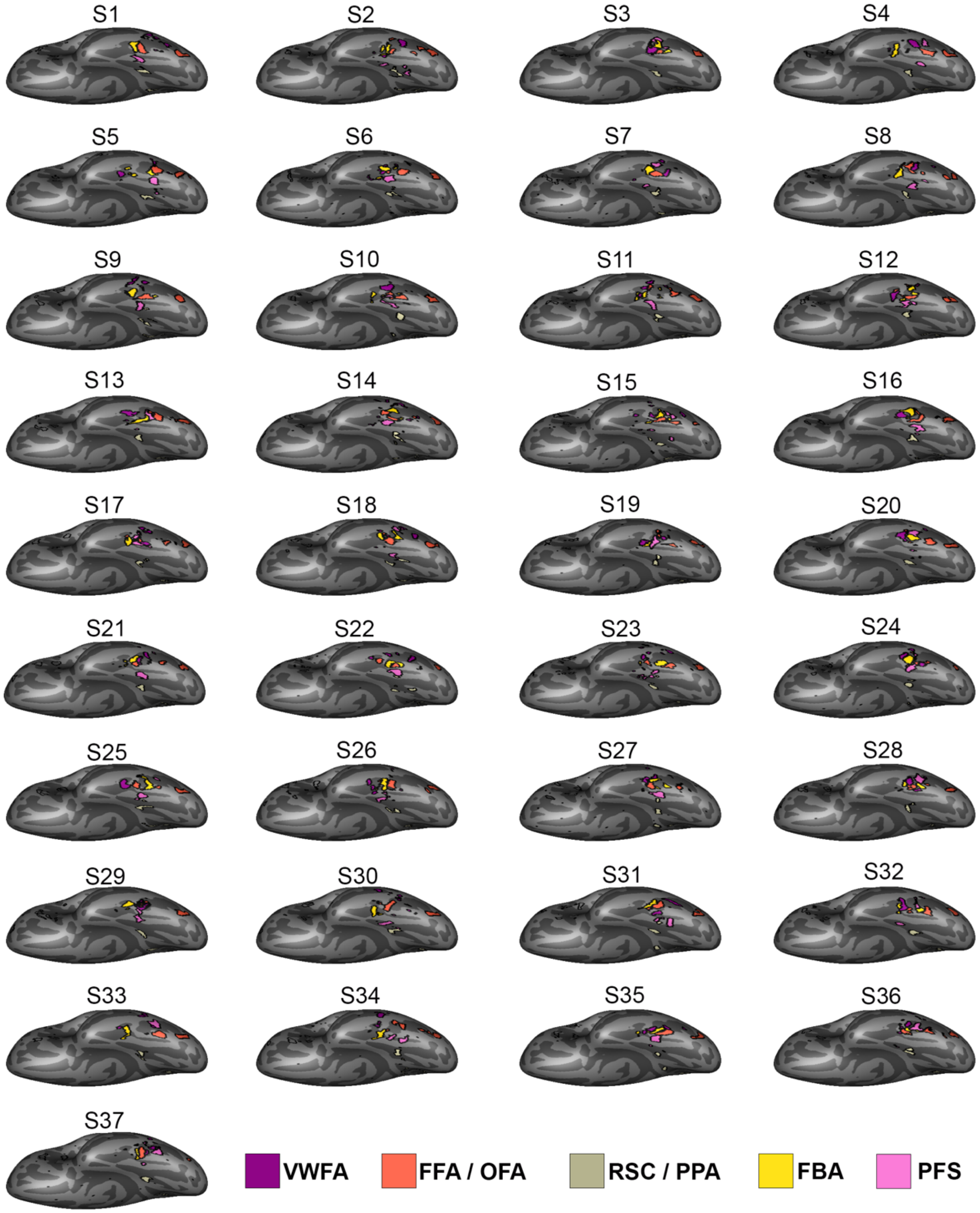

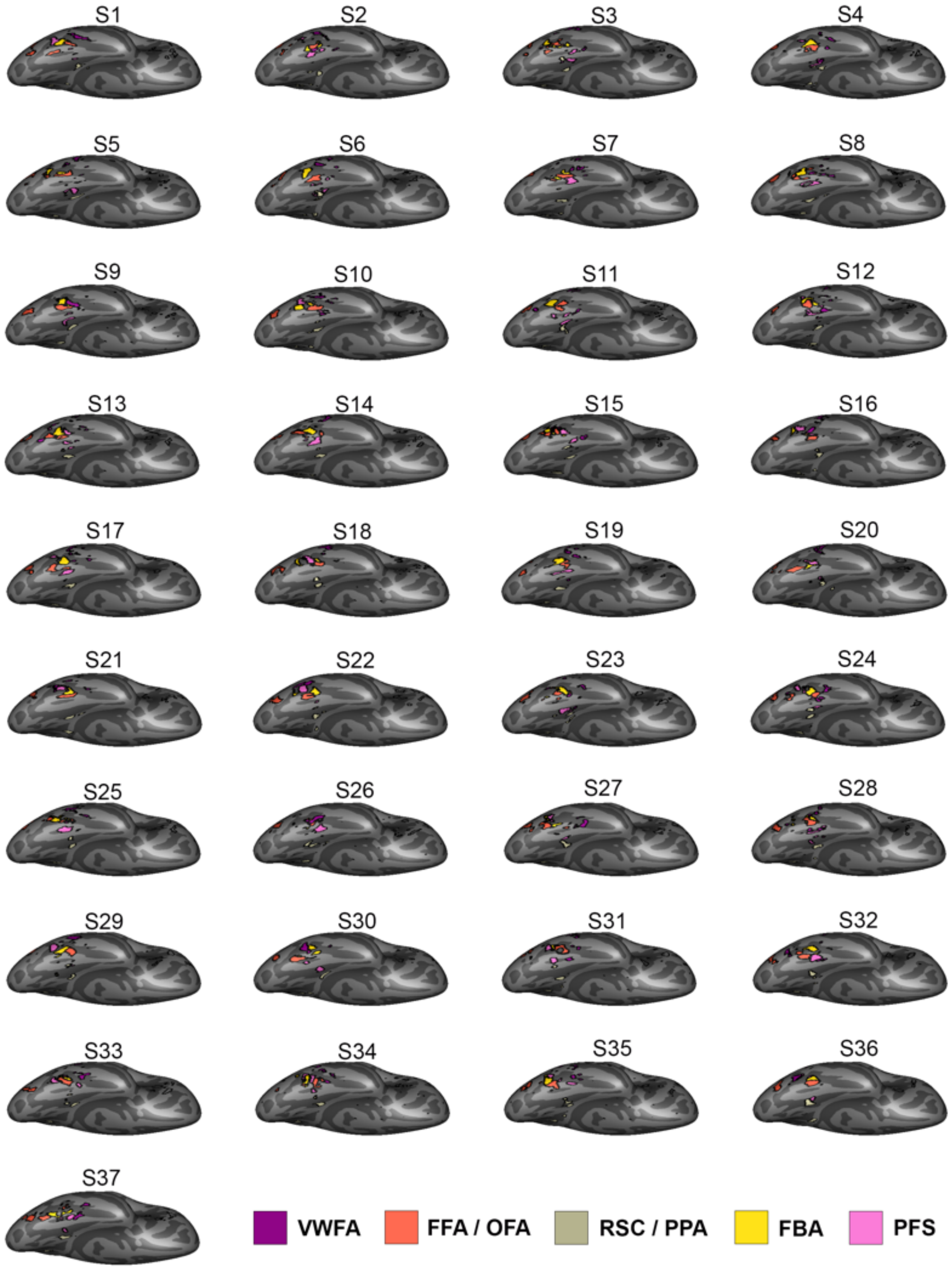
S1
S2
S3
S4
S5
S6
S7
S8
S9
S10
S11
S12
S13
S14
S15
S16
S17
S18
S19
S20
S21
S22
S23
S24
S25
S26
S27
S28
S29
S30
S31
S32
S33
S34
S35
S36
S37
VWFA
FFA / OFA
RSC / PPA
FBA
PFS

**Figure S2. rVWFA functional response profile**. A. VTC fROIs in the right hemisphere for an example subject. B. Functional profile of the right VWFA. The mean percent signal change (with standard error bars) to various visual and non-visual conditions are plotted. The preferred category, words, has a thick black outline. Individual subject PSCs are shown with grey circles. Significance is noted (*p<0.05, Bonferroni-holm corrected for 13 comparisons) for the preferred category only, with a black lining showing all conditions significantly lower than the preferred category (see **Table S2** for details). C. Averaged time-course of VWFA's responses to blocks of different experimental conditions. Dark line for mean across all subjects and shading for standard error.

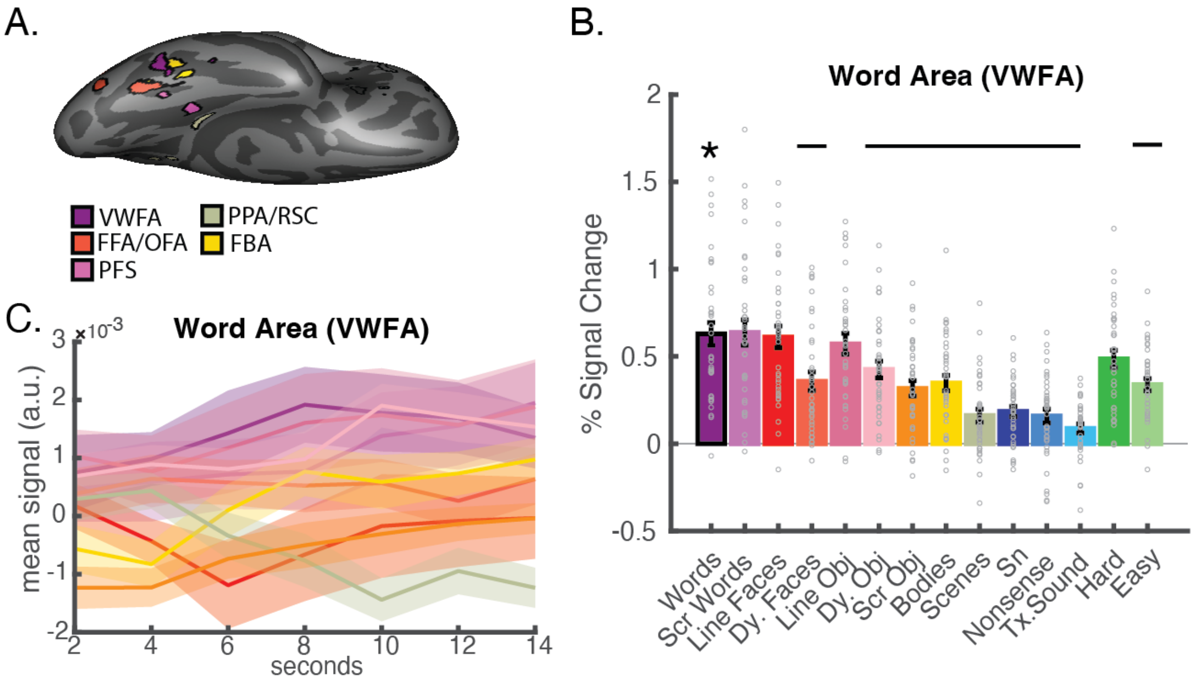

**Figure S3. Functional response profiles for right VTC category-selective fROIs. A.** The mean percent signal change (with standard error bars) to various visual and non-visual conditions are plotted. The preferred condition(s) for each fROI is outlined in black. Individual subject PSCs are shown with grey hollow circles. Significance is noted (*p<0.05, Bonferroni-holm corrected for 13 comparisons) for the preferred category only, with a black lining showing all conditions significantly lower than the preferred category (see **Table S2** for details). **B.** Averaged time-course of all VTC regions' responses to blocks of different experimental conditions. Dark line for mean across all subjects and shading for standard error. The same color scheme for denoting conditions as Figure S1 and Figure 1 is implemented for both A and B.

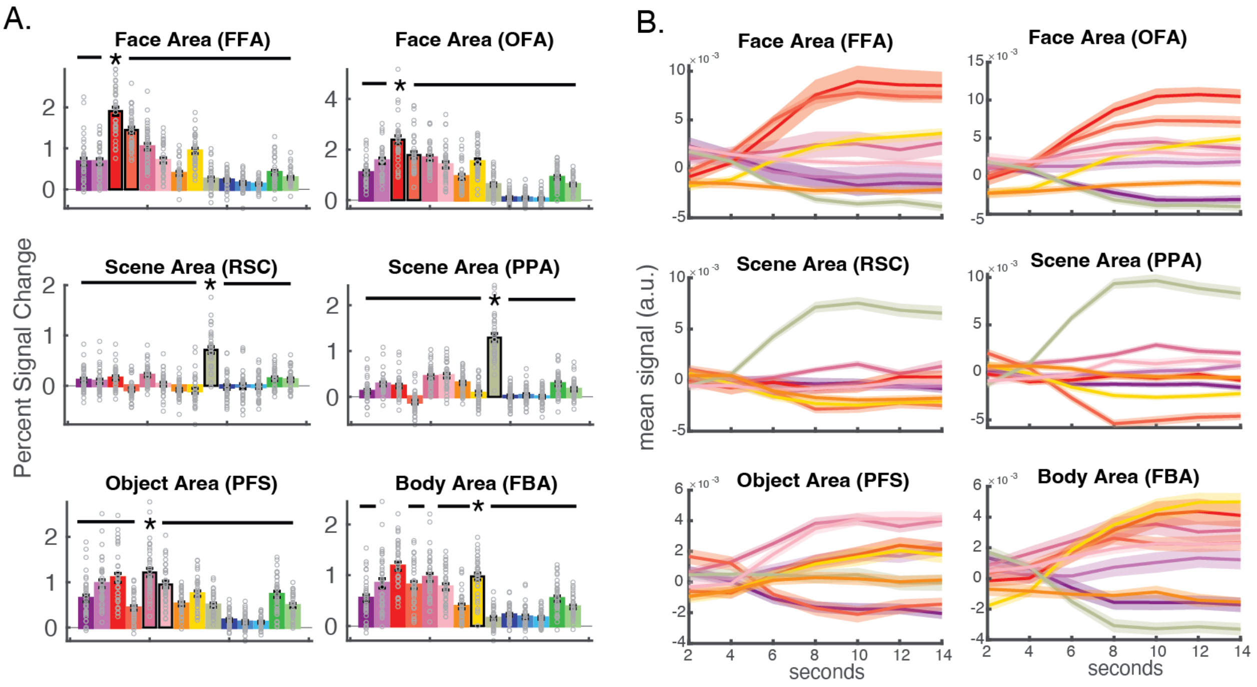

### Table S1. Comparing each fROI's preferred condition vs. other conditions for all methods of defining fROIs (LH).

Comparing each fROIs PSC to its preferred condition vs. other conditions (**fROI defined with the top 150 most significant voxels**, LH)

| Comparison Condition | Words | | Scrambled Words | | Line Faces | | Dynamic Faces | | Line Objects | | Dynamic Objects | | Scrambled Objects | | Bodies | | Scenes | |
|---|---|---|---|---|---|---|---|---|---|---|---|---|---|---|---|---|---|---|
| fROI (preferred condition) | t(df) | p | t(df) | p | t(df) | p | t(df) | p | t(df) | p | t(df) | p | t(df) | p | t(df) | p | t(df) | p |
| VWFA (words) | | | 7.03 (35) | $3.55x10^{-8}$ ** | 7.93 (35) | $2.50x10^{-9}$ ** | 8.73 (35) | $2.60x10^{-10}$ ** | 5.57 (35) | $2.87x10^{-6}$ ** | 7.19 (35) | $2.15x10^{-8}$ ** | 9.29 (35) | $5.65x10^{-11}$ ** | 7.91 (35) | $2.70x10^{-9}$ ** | 10.27 (35) | $4.20x10^{-12}$ ** |
| FFA (Faces) | 6.32 (36) | $2.59x10^{-7}$ ** | 10.02 (36) | $5.85x10^{-12}$ ** | | | 4.01 (35) | $3.01x10^{-4}$ ** | 8.26 (36) | $7.90x10^{-10}$ ** | 11.65 (35) | $1.34x10^{-13}$ ** | 12.68 (35) | $1.21x10^{-14}$ ** | 8.67 (35) | $3.05x10^{-10}$ ** | 14.52 (36) | $1.28x10^{-16}$ ** |
| OFA (Faces) | 8.23 (34) | $1.32x10^{-9}$ ** | 8.52 (34) | $6.04x10^{-10}$ ** | | | 4.71 (33) | $4.29x10^{-5}$ ** | 4.87 (33) | $2.73x10^{-5}$ ** | 5.19 (33) | $1.06x10^{-5}$ ** | 7.04 (33) | $4.70x10^{-8}$ ** | 6.62 (34) | $1.37x10^{-7}$ ** | 9.44 (34) | $5.02x10^{-11}$ ** |
| RSC (Scenes) | 5.67 (34) | $2.31x10^{-6}$ ** | 6.17 (34) | $5.17x10^{-7}$ ** | 5.47 (34) | $4.26x10^{-6}$ ** | 9.18 (35) | $7.48x10^{-11}$ ** | 5.12 (33) | $1.28x10^{-5}$ ** | 11.44 (35) | $2.23x10^{-13}$ ** | 10.6 (35) | $1.82x10^{-12}$ ** | 12.2 (35) | $3.63x10^{-14}$ ** | | |
| PPA (Scenes) | 11.27 (36) | $2.30x10^{-13}$ ** | 9.42 (36) | $3.02x10^{-11}$ ** | 10.41 (36) | $2.08x10^{-12}$ ** | 14.01 (36) | $3.80x10^{-16}$ ** | 7.86 (36) | $2.52x10^{-9}$ ** | 10.54 (36) | $1.48x10^{-12}$ ** | 11.97 (36) | $4.15x10^{-14}$ ** | 13.5 (36) | $1.18x10^{-15}$ ** | | |
| PFS (Objects) | 8.79 (35) | $2.21x10^{-10}$ ** | 6.57 (34) | $1.58x10^{-7}$ ** | 7.12 (34) | $3.11x10^{-8}$ ** | 8.99 (35) | $1.27x10^{-10}$ ** | | | 1.77 (35) | $8.00x10^{-2}$ | 6.58 (35) | $1.35x10^{-7}$ ** | 5.27 (35) | $7.17x10^{-6}$ ** | 5.87 (35) | $1.14x10^{-6}$ ** |
| FBA (Bodies) | 1.24 (36) | $2.20x10^{-1}$ | 4.2 (36) | $1.70x10^{-4}$ ** | -0.85 (36) | $4.00x10^{-1}$ | 3.9 (35) | $4.22x10^{-4}$ ** | -0.16 (36) | $8.80x10^{-1}$ | 3.54 (36) | $1.13x10^{-3}$ ** | 11.05 (35) | $5.88x10^{-13}$ ** | | | 9.95 (35) | $9.76x10^{-12}$ ** |

Comparing each fROIs PSC to its preferred condition vs. other conditions (**fROI defined with top 10% voxels**, LH)

| Comparison Condition | Words | | Scrambled Words | | Line Faces | | Dynamic Faces | | Line Objects | | Dynamic Objects | | Scrambled Objects | | Bodies | | Scenes | |
|---|---|---|---|---|---|---|---|---|---|---|---|---|---|---|---|---|---|---|
| fROI (preferred condition) | t(df) | p | t(df) | p | t(df) | p | t(df) | p | t(df) | p | t(df) | p | t(df) | p | t(df) | p | t(df) | p |
| VWFA (words) | | | 7.02 (36) | $3.08x10^{-8}$ ** | 7.26 (35) | $1.78x10^{-8}$ ** | 7.98 (36) | $1.80x10^{-9}$ ** | 5.27 (36) | $6.61x10^{-6}$ ** | 6.52 (36) | $1.41x10^{-7}$ ** | 9 (35) | $1.23x10^{-10}$ ** | 6.93 (36) | $4.03x10^{-8}$ ** | 9.59 (36) | $1.87x10^{-11}$ ** |
| FFA (Faces) | 6.67 (36) | $8.81x10^{-8}$ ** | 10.47 (36) | $1.80x10^{-12}$ ** | | | 4.07 (35) | $2.53x10^{-4}$ ** | 8.39 (36) | $5.41x10^{-10}$ ** | 12.15 (35) | $4.07x10^{-14}$ ** | 13.11 (36) | $2.85x10^{-15}$ ** | 9.03 (35) | $1.13x10^{-10}$ ** | 14.69 (35) | $1.56x10^{-16}$ ** |
| OFA (Faces) | 9.42 (30) | $1.79x10^{-10}$ ** | 8.27 (31) | $2.44x10^{-9}$ ** | | | 4.41 (31) | $1.15x10^{-4}$ ** | 5.3 (30) | $1.00x10^{-5}$ ** | 6.53 (31) | $2.77x10^{-7}$ ** | 8.74 (31) | $7.19x10^{-10}$ ** | 6.5 (31) | $2.98x10^{-7}$ ** | 10.21 (31) | $1.94x10^{-11}$ ** |
| RSC (Scenes) | 5.79 (35) | $1.46x10^{-6}$ ** | 6.39 (35) | $2.35x10^{-7}$ ** | 5.59 (35) | $2.71x10^{-6}$ ** | 9.08 (36) | $7.64x10^{-11}$ ** | 5.1 (34) | $1.27x10^{-5}$ ** | 11.06 (36) | $3.90x10^{-13}$ ** | 10.87 (36) | $6.35x10^{-13}$ ** | 12.33 (36) | $1.73x10^{-14}$ ** | | |
| PPA (Scenes) | 11.69 (36) | $8.18x10^{-14}$ ** | 9.96 (36) | $6.84x10^{-12}$ ** | 11.04 (36) | $4.18x10^{-13}$ ** | 14.72 (36) | $8.27x10^{-17}$ ** | 8.17 (36) | $1.01x10^{-9}$ ** | 11.03 (36) | $4.22x10^{-13}$ ** | 13.29 (36) | $1.90x10^{-15}$ ** | 14.05 (36) | $3.47x10^{-16}$ ** | | |
| PFS (Objects) | 9.04 (35) | $1.12x10^{-10}$ ** | 6.71 (34) | $1.04x10^{-7}$ ** | 7.24 (34) | $2.21x10^{-8}$ ** | 9.36 (35) | $4.63x10^{-11}$ ** | | | 2 (35) | $5.00x10^{-2}$ | 6.77 (35) | $7.55x10^{-8}$ ** | 5.61 (35) | $2.52x10^{-6}$ ** | 6.1 (35) | $5.74x10^{-7}$ ** |
| FBA (Bodies) | 1.26 (36) | $2.20x10^{-1}$ | 4.06 (36) | $2.50x10^{-4}$ ** | -0.85 (36) | $4.00x10^{-1}$ | 3.82 (35) | $5.19x10^{-4}$ ** | -0.23 (36) | $8.20x10^{-1}$ | 3.98 (36) | $3.21x10^{-4}$ ** | 11.28 (35) | $3.32x10^{-13}$ ** | | | 10.16 (35) | $5.61x10^{-12}$ ** |

Comparing each fROIs PSC to its preferred condition vs. other conditions (**fROI defined with a hard threshold (p<0.005)**, LH)

| Comparison Condition | Words | | Scrambled Words | | Line Faces | | Dynamic Faces | | Line Objects | | Dynamic Objects | | Scrambled Objects | | Bodies | | Scenes | |
|---|---|---|---|---|---|---|---|---|---|---|---|---|---|---|---|---|---|---|
| fROI (preferred condition) | t(df) | p | t(df) | p | t(df) | p | t(df) | p | t(df) | p | t(df) | p | t(df) | p | t(df) | p | t(df) | p |
| VWFA (words) | | | 6.46 (34) | $2.17x10^{-7}$ ** | 8.21 (34) | $1.41x10^{-9}$ ** | 11.35 (33) | $6.22x10^{-13}$ ** | 6.25 (34) | $4.09x10^{-7}$ ** | 8.67 (34) | $3.97x10^{-10}$ ** | 9.61 (34) | $3.20x10^{-11}$ ** | 9.43 (34) | $5.15x10^{-11}$ ** | 10.28 (34) | $5.75x10^{-12}$ ** |
| FFA (Faces) | 6.88 (35) | $5.49x10^{-8}$ ** | 11.49 (35) | $1.97x10^{-13}$ ** | | | 4.07 (35) | $2.53x10^{-4}$ ** | 8.86 (35) | $1.82x10^{-10}$ ** | 10.76 (35) | $1.22x10^{-12}$ ** | 11.49 (34) | $2.96x10^{-13}$ ** | 8.04 (35) | $1.86x10^{-9}$ ** | 13.56 (34) | $2.81x10^{-15}$ ** |
| OFA (Faces) | 8.53 (19) | $6.40x10^{-8}$ ** | 7.8 (19) | $2.45x10^{-7}$ ** | | | 2.91 (19) | $9.05x10^{-3}$ ** | 7.07 (19) | $1.00x10^{-6}$ ** | 4.76 (19) | $1.36x10^{-4}$ ** | 6.35 (19) | $4.30x10^{-6}$ ** | 5.39 (19) | $3.36x10^{-5}$ ** | 8.27 (19) | $1.02x10^{-7}$ ** |
| RSC (Scenes) | 6.11 (34) | $6.12x10^{-7}$ ** | 7.08 (34) | $3.57x10^{-8}$ ** | 5.5 (34) | $3.90x10^{-6}$ ** | 6.03 (36) | $6.31x10^{-7}$ ** | 4.08 (35) | $2.49x10^{-4}$ ** | 7.16 (36) | $2.00x10^{-8}$ ** | 8.59 (36) | $3.04x10^{-10}$ ** | 8.42 (36) | $4.95x10^{-10}$ ** | | |
| PPA (Scenes) | 8.26 (36) | $7.88x10^{-10}$ ** | 7.32 (36) | $1.27x10^{-8}$ ** | 7.67 (36) | $4.49x10^{-9}$ ** | 10.95 (36) | $5.21x10^{-13}$ ** | 5.39 (36) | $4.54x10^{-6}$ ** | 7.48 (35) | $9.39x10^{-9}$ ** | 10.65 (36) | $1.13x10^{-12}$ ** | 11.18 (36) | $2.89x10^{-13}$ ** | | |
| PFS (Objects) | 8.28 (33) | $1.45x10^{-9}$ ** | 4.78 (34) | $3.31x10^{-5}$ ** | 6.13 (34) | $5.86x10^{-7}$ ** | 10.63 (33) | $3.43x10^{-12}$ ** | | | 1.88 (34) | $7.00x10^{-2}$ | 4.71 (34) | $4.12x10^{-5}$ ** | 4.69 (34) | $4.30x10^{-5}$ ** | 5.23 (34) | $8.68x10^{-6}$ ** |
| FBA (Bodies) | 0.3 (33) | $7.60x10^{-1}$ | 1.86 (33) | $7.00x10^{-2}$ | -1.12 (33) | $2.70x10^{-1}$ | 3.31 (32) | $2.29x10^{-3}$ ** | -1.01 (33) | $3.20x10^{-1}$ | 2.47 (33) | $1.89x10^{-2}$ * | 9.42 (32) | $9.67x10^{-11}$ ** | | | 9.03 (33) | $1.97x10^{-10}$ ** |

### Table S2. Comparing each fROI's preferred condition vs. other conditions for all methods of defining fROIs (RH).

Comparing each fROIs PSC to its preferred condition vs. other conditions (**fROI defined with the top 150 most significant voxels**, RH)

| Comparison Condition | Words | | Scrambled Words | | Line Faces | | Dynamic Faces | | Line Objects | | Dynamic Objects | | Scrambled Objects | | Bodies | | Scenes | |
|---|---|---|---|---|---|---|---|---|---|---|---|---|---|---|---|---|---|---|
| fROI (preferred condition) | t(df) | p | t(df) | p | t(df) | p | t(df) | p | t(df) | p | t(df) | p | t(df) | p | t(df) | p | t(df) | p |
| VWFA (words) | | | -0.2 (36) | 8.40x10$^{-1}$ | 0.41 (36) | 6.80x10$^{-1}$ | 3.94 (36) | 3.61x10$^{-4}$ ** | 1.34 (35) | 1.90x10$^{-1}$ | 3.25 (36) | 2.49x10$^{-3}$ ** | 4.67 (36) | 4.07x10$^{-5}$ ** | 4.26 (36) | 1.41x10$^{-4}$ ** | 6.82 (36) | 5.68x10$^{-8}$ ** |
| FFA (Faces) | 10.59 (36) | 1.31x10$^{-12}$ ** | 14.18 (35) | 4.46x10$^{-16}$ ** | | | 4.4 (36) | 9.32x10$^{-5}$ ** | 8.48 (36) | 4.23x10$^{-10}$ ** | 11.71 (36) | 7.83x10$^{-14}$ ** | 14.07 (36) | 3.32x10$^{-16}$ ** | 9.37 (36) | 3.39x10$^{-11}$ ** | 14.11 (36) | 3.07x10$^{-16}$ ** |
| OFA (Faces) | 12.65 (32) | 5.44x10$^{-14}$ ** | 9.17 (32) | 1.82x10$^{-10}$ ** | | | 5.72 (32) | 2.41x10$^{-6}$ ** | 6.81 (32) | 1.07x10$^{-7}$ ** | 6.12 (32) | 7.68x10$^{-7}$ ** | 9.28 (32) | 1.36x10$^{-10}$ ** | 5.67 (32) | 2.83x10$^{-6}$ ** | 11.06 (32) | 1.81x10$^{-12}$ ** |
| RSC (Scenes) | 7.32 (34) | 1.78x10$^{-8}$ ** | 7.21 (33) | 2.86x10$^{-8}$ ** | 7.26 (33) | 2.48x10$^{-8}$ ** | 10.88 (35) | 8.97x10$^{-13}$ ** | 6.96 (33) | 5.93x10$^{-8}$ ** | 9.97 (35) | 9.06x10$^{-12}$ ** | 12.17 (35) | 3.95x10$^{-14}$ ** | 11.84 (35) | 8.48x10$^{-14}$ ** | | |
| PPA (Scenes) | 11.35 (36) | 1.89x10$^{-13}$ ** | 9.9 (35) | 1.09x10$^{-11}$ ** | 10.74 (36) | 8.98x10$^{-13}$ ** | 15.2 (36) | 3.03x10$^{-17}$ ** | 8.04 (35) | 1.84x10$^{-9}$ ** | 9.74 (36) | 1.24x10$^{-11}$ ** | 12.14 (36) | 2.72x10$^{-14}$ ** | 16.44 (35) | 4.88x10$^{-18}$ ** | | |
| PFS (Objects) | 11.22 (36) | 2.65x10$^{-13}$ ** | 4.32 (35) | 1.22x10$^{-4}$ ** | 2.18 (36) | 3.59x10$^{-2}$ ** | 8.06 (36) | 1.41x10$^{-9}$ ** | | | 3.03 (36) | 4.51x10$^{-3}$ ** | 7.72 (35) | 4.59x10$^{-9}$ ** | 4.91 (36) | 1.97x10$^{-5}$ ** | 7 (36) | 3.26x10$^{-8}$ ** |
| FBA (Bodies) | 4.8 (35) | 2.91x10$^{-5}$ ** | 1.9 (35) | 7.00x10$^{-2}$ | -1.99 (34) | 5.00x10$^{-2}$ | 3 (35) | 4.96x10$^{-3}$ ** | 0.37 (34) | 7.10x10$^{-1}$ | 3.88 (35) | 4.45x10$^{-4}$ ** | 10.6 (35) | 1.79x10$^{-12}$ ** | | | 11.52 (34) | 2.75x10$^{-13}$ ** |

Comparing each fROIs PSC to its preferred condition vs. other conditions (**fROI defined with top 10% voxels**, RH)

| Comparison Condition | Words | | Scrambled Words | | Line Faces | | Dynamic Faces | | Line Objects | | Dynamic Objects | | Scrambled Objects | | Bodies | | Scenes | |
|---|---|---|---|---|---|---|---|---|---|---|---|---|---|---|---|---|---|---|
| fROI (preferred condition) | t(df) | p | t(df) | p | t(df) | p | t(df) | p | t(df) | p | t(df) | p | t(df) | p | t(df) | p | t(df) | p |
| VWFA (words) | | | -1.06 (36) | 3.00x10$^{-1}$ | -0.7 (36) | 4.90x10$^{-1}$ | 3.35 (36) | 1.90x10$^{-3}$ ** | 0.73 (35) | 4.70x10$^{-1}$ | 2.07 (36) | 4.60x10$^{-2}$ * | 4.13 (36) | 2.07x10$^{-4}$ ** | 3.09 (36) | 3.85x10$^{-3}$ ** | 6.74 (36) | 7.20x10$^{-8}$ ** |
| FFA (Faces) | 12.08 (35) | 4.86x10$^{-14}$ ** | 13.97 (35) | 7.03x10$^{-16}$ ** | | | 3.84 (36) | 4.83x10$^{-4}$ ** | 8.46 (36) | 4.39x10$^{-10}$ ** | 10.89 (36) | 6.06x10$^{-13}$ ** | 13.23 (36) | 2.16x10$^{-15}$ ** | 8.69 (36) | 2.30x10$^{-10}$ ** | 13.42 (36) | 1.41x10$^{-15}$ ** |
| OFA (Faces) | 12.28 (32) | 1.18x10$^{-13}$ ** | 8.69 (32) | 6.29x10$^{-10}$ ** | | | 5.07 (32) | 1.61x10$^{-5}$ ** | 6.72 (32) | 1.37x10$^{-7}$ ** | 6.31 (32) | 4.39x10$^{-7}$ ** | 9.41 (32) | 9.78x10$^{-11}$ ** | 5.77 (32) | 2.10x10$^{-6}$ ** | 11.44 (32) | 7.58x10$^{-13}$ ** |
| RSC (Scenes) | 8.25 (33) | 1.58x10$^{-9}$ ** | 7.61 (33) | 9.33x10$^{-9}$ ** | 7.55 (33) | 1.09x10$^{-8}$ ** | 11.26 (35) | 3.52x10$^{-13}$ ** | 7.11 (33) | 3.85x10$^{-8}$ ** | 10.53 (35) | 2.16x10$^{-12}$ ** | 13.33 (35) | 2.80x10$^{-15}$ ** | 12.67 (35) | 1.25x10$^{-14}$ ** | | |
| PPA (Scenes) | 11.77 (35) | 1.01x10$^{-13}$ ** | 10.23 (35) | 4.63x10$^{-12}$ ** | 11.16 (35) | 4.47x10$^{-13}$ ** | 15.59 (36) | 1.38x10$^{-17}$ ** | 8.22 (35) | 1.09x10$^{-9}$ ** | 10 (36) | 6.26x10$^{-12}$ ** | 12.42 (36) | 1.40x10$^{-14}$ ** | 17.36 (35) | 8.98x10$^{-19}$ ** | | |
| PFS (Objects) | 11.16 (36) | 3.02x10$^{-13}$ ** | 4.25 (35) | 1.52x10$^{-4}$ ** | 2.1 (36) | 4.31x10$^{-2}$ ** | 7.81 (36) | 2.91x10$^{-9}$ ** | | | 2.92 (36) | 6.00x10$^{-3}$ ** | 7.71 (35) | 4.74x10$^{-9}$ ** | 4.68 (36) | 4.03x10$^{-5}$ ** | 6.8 (36) | 6.09x10$^{-8}$ ** |
| FBA (Bodies) | 6.13 (35) | 5.15x10$^{-7}$ ** | 3.13 (35) | 3.48x10$^{-3}$ ** | -1.81 (35) | 8.00x10$^{-2}$ | 3.09 (35) | 3.92x10$^{-3}$ ** | 0.76 (35) | 4.50x10$^{-1}$ | 4.53 (35) | 6.49x10$^{-5}$ ** | 12.56 (35) | 1.60x10$^{-14}$ ** | | | 13.48 (34) | 3.39x10$^{-15}$ ** |

Comparing each fROIs PSC to its preferred condition vs. other conditions (**fROI defined with a hard threshold (p<0.005)**, RH)

| Comparison Condition | Words | | Scrambled Words | | Line Faces | | Dynamic Faces | | Line Objects | | Dynamic Objects | | Scrambled Objects | | Bodies | | Scenes | |
|---|---|---|---|---|---|---|---|---|---|---|---|---|---|---|---|---|---|---|
| fROI (preferred condition) | t(df) | p | t(df) | p | t(df) | p | t(df) | p | t(df) | p | t(df) | p | t(df) | p | t(df) | p | t(df) | p |
| VWFA (words) | | | -0.13 (28) | 9.00x10$^{-1}$ | -0.25 (27) | 8.00x10$^{-1}$ | 2.99 (28) | 5.72x10$^{-3}$ ** | 0.33 (28) | 7.40x10$^{-1}$ | 1.8 (28) | 8.00x10$^{-2}$ | 3.7 (27) | 9.74x10$^{-4}$ ** | 3.66 (27) | 1.08x10$^{-3}$ ** | 4.76 (28) | 5.28x10$^{-5}$ ** |
| FFA (Faces) | 10.01 (35) | 8.22x10$^{-12}$ ** | 13.92 (34) | 1.33x10$^{-15}$ ** | | | 4.72 (34) | 3.98x10$^{-5}$ ** | 11.27 (34) | 5.04x10$^{-13}$ ** | 10.2 (35) | 4.97x10$^{-12}$ ** | 11.23 (35) | 3.79x10$^{-13}$ ** | 7.15 (35) | 2.43x10$^{-8}$ ** | 13.11 (34) | 7.43x10$^{-15}$ ** |
| OFA (Faces) | 8.8 (22) | 1.17x10$^{-8}$ ** | 9.08 (22) | 6.73x10$^{-9}$ ** | | | 3.9 (22) | 7.73x10$^{-4}$ ** | 6.63 (22) | 1.16x10$^{-6}$ ** | 6.18 (22) | 3.19x10$^{-6}$ ** | 11.6 (22) | 7.59x10$^{-11}$ ** | 5.49 (22) | 1.62x10$^{-5}$ ** | 11.38 (22) | 1.10x10$^{-10}$ ** |
| RSC (Scenes) | 6.57 (35) | 1.38x10$^{-7}$ ** | 5.18 (36) | 8.53x10$^{-6}$ ** | 4.18 (36) | 1.77x10$^{-4}$ ** | 6.43 (36) | 1.83x10$^{-7}$ ** | 4.24 (36) | 1.48x10$^{-4}$ ** | 7.44 (36) | 8.80x10$^{-9}$ ** | 10.17 (36) | 3.93x10$^{-12}$ ** | 9.7 (36) | 1.39x10$^{-11}$ ** | | |
| PPA (Scenes) | 10.33 (36) | 2.61x10$^{-12}$ ** | 8.86 (35) | 1.83x10$^{-10}$ ** | 11.84 (35) | 8.62x10$^{-14}$ ** | 16.13 (36) | 4.68x10$^{-18}$ ** | 6.16 (36) | 4.26x10$^{-7}$ ** | 8.95 (36) | 1.10x10$^{-10}$ ** | 12.46 (35) | 2.01x10$^{-14}$ ** | 14.35 (36) | 1.83x10$^{-16}$ ** | | |
| PFS (Objects) | 7.74 (31) | 9.88x10$^{-9}$ ** | 4.59 (31) | 6.96x10$^{-5}$ ** | 0.57 (31) | 5.70x10$^{-1}$ | 8.09 (30) | 5.01x10$^{-9}$ ** | | | 3.3 (31) | 2.43x10$^{-3}$ ** | 7.68 (31) | 1.17x10$^{-8}$ ** | 5.93 (30) | 1.69x10$^{-6}$ ** | 8.85 (31) | 5.45x10$^{-10}$ ** |
| FBA (Bodies) | 4.96 (31) | 2.44x10$^{-5}$ ** | 1.52 (32) | 1.40x10$^{-1}$ | -0.78 (31) | 4.40x10$^{-1}$ | 2.82 (32) | 8.10x10$^{-3}$ ** | -0.18 (32) | 8.50x10$^{-1}$ | 2.27 (32) | 3.03x10$^{-2}$ * | 8.35 (32) | 1.55x10$^{-9}$ ** | | | 9.3 (32) | 1.31x10$^{-10}$ ** |

**Table S3. Comparing category selectivity among left VTC**

| Word Selectivity | | |
|---|---|---|
| VWFA vs. | t(df) | p |
| FBA | 7.35(31) | 8.61e-08** |
| FFA | 5.13(31) | 1.47e-05** |
| OFA | 8.91(31) | 2.32e-09** |
| PFS | 7.84(31) | 3.044e-08** |
| PPA | 9.77(31) | 3.306e-10** |
| RSC | 6.52(31) | 5.64e-07** |

| Body Selectivity | | |
|---|---|---|
| FBA vs. | t(df) | p |
| FFA | 2.63(35) | 0.013** |
| OFA | 3.66(35) | 0.00237** |
| PFS | 3.68(35) | 0.00237** |
| PPA | 5.72(35) | 8.95e-06** |
| RSC | 5.72(35) | 8.95e-06** |
| VWFA | 6.03(35) | 4.272e-06** |

| Face Selectivity | | |
|---|---|---|
| FFA vs. | t(df) | p |
| FBA | 9.61(36) | 5.4e-11** |
| OFA | 4.26(36) | 0.000138** |
| PFS | 12.65(36) | 4.095e-14** |
| PPA | 18.24(36) | 5.424e-19** |
| RSC | 11.91(36) | 1.924e-13** |
| VWFA | 8.6(36) | 5.98e-10** |

| Scene Selectivity | | |
|---|---|---|
| PPA vs. | t(df) | p |
| FBA | 15.95(34) | 1.428e-16** |
| FFA | 15.57(34) | 1.94e-16** |
| OFA | 15.71(34) | 1.87e-16** |
| PFS | 10.56(34) | 5.74e-12** |
| RSC | 0.21(34) | 8.31E-01 |
| VWFA | 14.86(34) | 5.88e-16** |

| Face Selectivity | | |
|---|---|---|
| OFA vs. | t(df) | p |
| FBA | 1.22(36) | 2.32E-01 |
| FFA | -4.26(36) | 5.52e-04** |
| PFS | 5.7(36) | 8.75e-06** |
| PPA | 8.24(36) | 4.974e-09** |
| RSC | 4.16(36) | 5.58e2-4** |
| VWFA | 2.95(36) | 0.01** |

| Scene Selectivity | | |
|---|---|---|
| RSC vs. | t(df) | p |
| FBA | 11.85(34) | 7.68e-13** |
| FFA | 11.16(34) | 2.596e-12** |
| OFA | 10.81(34) | 4.65e-12** |
| PFS | 8.15(34) | 3.36e-09** |
| PPA | -0.21(34) | 8.31E-01 |
| VWFA | 11.33(34) | 2.155e-12** |

| Object Selectivity | | |
|---|---|---|
| PFS vs. | t(df) | p |
| FBA | 7.69(36) | 1.656e-08** |
| FFA | 10.27(36) | 1.53e-11** |
| OFA | 10.75(36) | 5.256e-12** |
| PPA | 3.88(36) | 0.000424** |
| RSC | 6.69(36) | 2.541e-07** |
| VWFA | 5.53(36) | 5.9e-06** |

**Table S4. Language-selective effect and attentional effect in the left VTC fROIs**

**Language-selective effect in the left VTC fROIs**

| fROIs | Contrast | t(df) | p |
|---|---|---|---|
| FFA | Sent > Ns | t(35)=1.93 | 0.061 |
| OFA | Sent > Ns | t(35)=0.81 | 0.425 |
| PFS | Sent > Ns | t(35)=1.06 | 0.298 |
| PPA | Sent > Ns | t(36)=-0.5 | 0.623 |
| RSC | Sent > Ns | t(36)=-0.39 | 0.701 |
| VWFA | Sent > Ns | t(35)=2.85 | 0.007** |

**Attentiona effect in the left VTC fROIs**

| fROIs | Contrast | t(df) | p |
|---|---|---|---|
| FFA | Hard > Easy | t(36)=4.77 | 3.02E-05** |
| OFA | Hard > Easy | t(36)=6.75 | 6.98E-08** |
| PFS | Hard > Easy | t(36)=7.74 | 3.56E-09** |
| PPA | Hard > Easy | t(36)=5.94 | 8.29E-07** |
| RSC | Hard > Easy | t(35)=-0.38 | 7.04E-01 |
| VWFA | Hard > Easy | t(36)=4.22 | 1.58E-04** |

**Supplementary Results: Replication for new VWFA parameters.**
Given that our static visual localizer was collected with different scanning parameters than the other three localizers, we collected additional data from eight participants to ensure our results were not driven by different scan parameters. Specifically, These subjects completed two more runs of the static visual localizer (in addition to two runs of all other tasks), with identical scan parameters for all tasks (TR=1000ms, TE=28ms, voxel resolution of 2x2x3 $mm^3$, 120 × 120 base resolution, 56 slices for the whole-brain coverage).
We replicate our main results, showing the left VWFA is still numerically more responsive to words than any other condition (**Table S5**), although not all differences are significant ($p<0.08$), likely due to small N). The other fROIs (FFA, OFA, RSC, PPA, and PFS) also show the highest responsiveness to their preferred categories (except for the FBA, which was excluded from the further analyses), typically reaching significance (**Table S5**). Overall, a functional profile trend similar to the main results is seen in even this small sample of subjects across left and right VTC fROIs (**see Figure S4**).

After confirming the category selectivity in the VWFA as well as other VTC fROIs, we then further examined the VWFA's response pattern to non-word visual as well as auditory language conditions based on the results observed in the main results. Planned paired t-tests were used and one-tailed uncorrected p values are reported.

As shown in **Figure S4** below, in addition to the absolute highest responses to the visual words, the VWFA responded the second highest to the object conditions: the averaged responses to the line-drawing objects and dynamic objects significantly is higher than the mean of other non-word conditions ($t(7)=1.86$, $p=0.053$). When compared to the scrambled objects in the dynamic localizer, the VWFA shows significantly higher responses to not only objects ($t(7)=4.72$, $p=0.001$) but also faces ($t(7)=2.53$, $p=0.02$, one-tailed) and bodies ($t(7)=2.03$, $p=0.041$).

Examining language selectivity in this smaller sample, again, we found that the VWFA is the only VTC region that shows significant higher response to Sn versus Ns ($t(7)=3.15$, $p=0.008$) AND for Sn versus Tx ($t(7)=1.93$, $p=0.048$) (**Table S5**, left). Similar to the main results, in contrast to the unique language sensitivity observed in the VWFA, the attentional effect was shown in almost all VTC regions except for the RSC (**Table S5**, right). We then compared the language-selective response (Sn > Ns) in the VWFA and canonical language regions with two-way rmANOVA. When comparing the VWFA to the temporal language regions, we found significant condition ($F(1,7)=38.79$, $p=4.33 \times 10^{-4}$) and fROI effects ($F(1,7)=35.61$, $p=5.60 \times 10^{-4}$), as well as interaction ($F(1,7)=23.44$, $p=0.002$), indicating that the language sensitivity is smaller in the VWFA; but the VWFA is not different from the frontal language regions (condition: $F(1,7)=10.11$, $p=0.016$; fROI: $F(1,7)=0.24$, $p=0.639$; interaction: $F(1,7)=2.66$, $p=0.147$)

In this sample, motion across tasks was not different (one-way repeated measures ANOVA examining mean framewise displacement (averaged across runs): $F(3,21)=0.733$, $p=0.54$). Additionally, the temporal signal to noise ratio (tSRN) was comparable across tasks in the lVWFA and for most comparisons of other regions. A two-way repeated measure ANOVA (task tSNR by fROI) showed a significant effect of task ($F(3,21)=10.29$, $p=2.26x10^{-4}$) and interaction between task and fROI ($F(15,105)=6.02$, $p=6.93x10^{-9}$). The follow-up t-tests showed a significant effect of task in all left VTC fROIs (all $p<0.05$ except the RSC, $p=0.32$); however, follow-up paired-samples t-tests across tasks with the lVWFA show no significant differences (all $t<3.7$ and all Bonferroni-holm $p>0.05$). Additionally, the lVWFA tSNR was always numerically lowest in the static visual (with matching parameters) localizer. Our results do not appear to be explainable by differences in scanning parameters, signal to noise ratio, or motion.

#### Table S5. Comparing each fROIs PSC to its preferred condition vs. other conditions (matching parameter subset)

fROIs defined with the top 150 most significant voxels (**LH**)

| Comparison Condition | Words | | Scrambled Words | | Line Faces | | Dynamic Faces | | Line Objects | | Dynamic Objects | | Scrambled Objects | | Bodies | | Scenes | |
|---|---|---|---|---|---|---|---|---|---|---|---|---|---|---|---|---|---|---|
| **fROI (preferred condition)** | t(df) | p | t(df) | p | t(df) | p | t(df) | p | t(df) | p | t(df) | p | t(df) | p | t(df) | p | t(df) | p |
| VWFA (words) | | | 2.2 (7) | 6.00x10$^{-2}$ | 2.23 (7) | 6.00x10$^{-2}$ | 2.63 (7) | 3.40x10$^{-2}$ * | 2.17 (7) | 7.00x10$^{-2}$ | 1.66 (7) | 1.40x10$^{-1}$ | 4.93 (7) | 1.70x10$^{-3}$ ** | 2.1 (7) | 7.00x10$^{-2}$ | 4.47 (7) | 2.91x10$^{-3}$ ** |
| FFA (Faces) | 4.14 (7) | 4.33x10$^{-3}$ ** | 4.79 (7) | 1.99x10$^{-3}$ ** | | | -1.6 (7) | 1.50x10$^{-1}$ | 4 (7) | 5.19x10$^{-3}$ ** | 2.79 (7) | 2.71x10$^{-2}$ * | 4.45 (7) | 2.96x10$^{-3}$ ** | 2.31 (7) | 5.00x10$^{-2}$ | 4.65 (7) | 2.33x10$^{-3}$ ** |
| OFA (Faces) | 3.7 (7) | 7.61x10$^{-3}$ * | 3.27 (7) | 1.37x10$^{-2}$ * | | | 1.77 (7) | 1.20x10$^{-1}$ | 2.47 (7) | 4.26x10$^{-2}$ * | 1.37 (7) | 2.10x10$^{-1}$ | 3.18 (7) | 1.55x10$^{-2}$ * | 2.13 (7) | 7.00x10$^{-2}$ | 4.33 (7) | 3.43x10$^{-3}$ ** |
| RSC (Scenes) | 3.93 (7) | 5.67x10$^{-3}$ ** | 3.89 (7) | 5.96x10$^{-3}$ ** | 4.01 (7) | 5.15x10$^{-3}$ ** | 2.89 (7) | 2.34x10$^{-2}$ ** | 3.18 (7) | 1.55x10$^{-2}$ ** | 7.23 (7) | 1.73x10$^{-4}$ ** | 3.96 (7) | 5.47x10$^{-3}$ ** | 5.55 (7) | 8.58x10$^{-4}$ ** | | |
| PPA (Scenes) | 6.3 (7) | 4.03x10$^{-4}$ ** | 4.62 (7) | 2.43x10$^{-3}$ ** | 5.8 (7) | 6.64x10$^{-4}$ ** | 5.53 (7) | 8.79x10$^{-4}$ ** | 4.08 (7) | 4.68x10$^{-3}$ ** | 3.89 (7) | 5.98x10$^{-3}$ ** | 4.88 (7) | 1.80x10$^{-3}$ ** | 5.98 (7) | 5.55x10$^{-4}$ ** | | |
| PFS (Objects) | 3.5 (7) | 9.98x10$^{-3}$ * | 2.21 (7) | 6.00x10$^{-2}$ | 5.32 (7) | 1.10x10$^{-3}$ ** | 4.04 (7) | 4.94x10$^{-3}$ ** | | | -1.98 (7) | 9.00x10$^{-2}$ | 2.24 (7) | 6.00x10$^{-2}$ | 1.37 (7) | 2.10x10$^{-1}$ | 2.34 (7) | 5.00x10$^{-2}$ |
| FBA (Bodies) | 3.65 (7) | 8.21x10$^{-3}$ ** | 5.41 (7) | 1.00x10$^{-3}$ ** | 1.15 (7) | 2.90x10$^{-1}$ | 0.68 (7) | 5.20x10$^{-1}$ | 4.65 (7) | 2.33x10$^{-3}$ ** | 2.25 (7) | 6.00x10$^{-2}$ | 7.58 (7) | 1.28x10$^{-4}$ ** | | | 6.64 (7) | 2.92x10$^{-4}$ ** |

fROIs defined with the top 150 most significant voxels (**RH**)

| Comparison Condition | Words | | Scrambled Words | | Line Faces | | Dynamic Faces | | Line Objects | | Dynamic Objects | | Scrambled Objects | | Bodies | | Scenes | |
|---|---|---|---|---|---|---|---|---|---|---|---|---|---|---|---|---|---|---|
| **fROI (preferred condition)** | t(df) | p | t(df) | p | t(df) | p | t(df) | p | t(df) | p | t(df) | p | t(df) | p | t(df) | p | t(df) | p |
| VWFA (words) | | | 0.84 (7) | 0.43 | 0.84 (7) | 0.43 | 1.53 (7) | 0.17 | 1.3 (7) | 0.23 | 0.14 (7) | 0.9 | 1.43 (7) | 0.2 | 0.7 (7) | 0.51 | 2.27 (7) | 0.06 |
| FFA (Faces) | 7.22 (7) | 1.75x10$^{-4}$ ** | 5.58 (7) | 8.35x10$^{-4}$ ** | | | -1.07 (7) | 0.32 | 5.03 (7) | 1.52x10$^{-3}$ ** | 2.95 (7) | 2.14x10$^{-2}$ ** | 5.79 (7) | 6.68x10$^{-4}$ ** | 3.27 (7) | 1.37x10$^{-2}$ ** | 7.53 (7) | 1.34x10$^{-4}$ ** |
| OFA (Faces) | 6.33 (7) | 3.93x10$^{-4}$ ** | 4.51 (7) | 2.76x10$^{-3}$ ** | | | 0.61 (7) | 0.56 | 4.3 (7) | 3.57x10$^{-3}$ ** | 3.72 (7) | 7.47x10$^{-3}$ ** | 6.1 (7) | 4.92x10$^{-4}$ ** | 4.69 (7) | 2.24x10$^{-3}$ ** | 6.02 (7) | 5.30x10$^{-4}$ ** |
| RSC (Scenes) | 7.28 (7) | 1.65x10$^{-4}$ ** | 5.8 (7) | 6.67 x10$^{-4}$ ** | 6.13 (7) | 4.76x10$^{-4}$ ** | 4.31 (7) | 3.53x10$^{-3}$ ** | 5.2 (7) | 1.25x10$^{-3}$ ** | 7.08 (7) | 1.98x10$^{-4}$ ** | 9.19 (7) | 3.7201.10$^{-5}$ * | 8.27 (7) | 7.3553x10$^{-5}$ ** | | |
| PPA (Scenes) | 8.19 (7) | 7.82x10$^{-5}$ ** | 6.3 (7) | 4.05x10$^{-4}$ ** | 7.46 (7) | 1.42x10$^{-4}$ ** | 8.61 (7) | 5.68x10$^{-5}$ ** | 5.36 (7) | 1.06x10$^{-3}$ ** | 3.86 (7) | 6.24x10$^{-3}$ ** | 5.29 (7) | 1.14x10$^{-3}$ ** | 7.03 (7) | 2.06x10$^{-4}$ ** | | |
| PFS (Objects) | 8.67 (7) | 5.42x10$^{-5}$ ** | 1.25 (7) | 0.25 | 1.64 (7) | 0.15 | 9.43 (7) | 3.14x10$^{-5}$ ** | | | -2.09 (7) | 0.08 | 3.13 (7) | 1.67x10$^{-2}$ * | -0.08 (7) | 0.94 | 3.6 (7) | 8.75x10$^{-3}$ * |
| FBA (Bodies) | 3.89 (7) | 5.95x10$^{-3}$ ** | 1.92 (7) | 0.1 | 0.65 (7) | 0.54 | 1.8 (7) | 0.11 | 1.93 (7) | 0.1 | 0.13 (7) | 0.9 | 4.92 (7) | 1.7x10$^{-3}$ ** | | | 4.54 (7) | 2.66x10$^{-3}$ ** |

**Table S6. Language- and attentional-selective responses in the left VTC fROIs.**

Language-selective effect in the VTC fROIs

| fROIs | Comparison | t(df) | p (uncorrected, one-tailed) |
|---|---|---|---|
| VWFA | Sent > Ns | 3.15(7) | 0.008* |
| VWFA | Sent >Tx | 1.93(7) | 0.048* |
| FFA | Sent > Ns | 2.52(7) | 0.020* |
| FFA | Sent >Tx | 1.37(7) | 0.107 |
| OFA | Sent > Ns | 1.5(7) | 0.089 |
| OFA | Sent >Tx | 0.31(7) | 0.383 |
| PFS | Sent > Ns | 3.14(7) | 0.008* |
| PFS | Sent >Tx | 0.99(7) | 0.179 |
| PPA | Sent > Ns | 2.59(7) | 0.018* |
| PPA | Sent >Tx | 0.54(7) | 0.305 |
| RSC | Sent > Ns | 2.26(7) | 0.029* |
| RSC | Sent >Tx | 0(7) | 0.498 |

Attentioal effect in the VTC fROIs

| fROIs | Comparison | t(df) | p (uncorrected, one-tailed) |
|---|---|---|---|
| VWFA | Hard > Easy | 2.86(7) | 0.012* |
| FFA | Hard > Easy | 2.13(7) | 0.036* |
| OFA | Hard > Easy | 3.69(7) | 0.004* |
| PFS | Hard > Easy | 3.76(7) | 0.004* |
| PPA | Hard > Easy | 3.73(7) | 0.004* |
| RSC | Hard > Easy | -0.03(7) | 0.4895 |

**Figure S4: Functional profile for left (A) and right (B) VTC fROIs in 8 subjects who completed the static visual localizer with scan parameters matching all other functional localizers.** Bars show mean across subjects with standard error bars. The expected preferred category is highlighted in black. Individual subject PSCs are shown with grey circles.

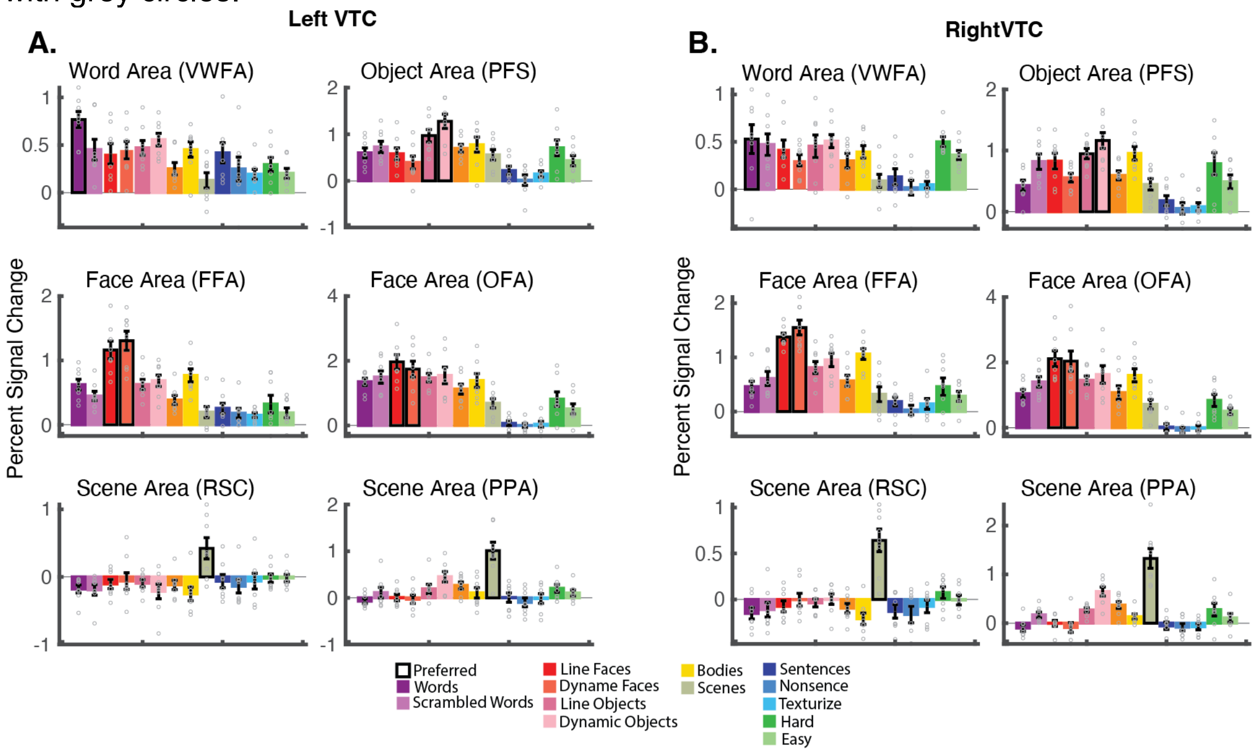

**Figure S5. Probabilistic maps for the language-selective response, defined with Sentences (Sn) > Nonsense Sounds (Ns), within the bilateral VTC.** Minimum overlap = 5 subjects.

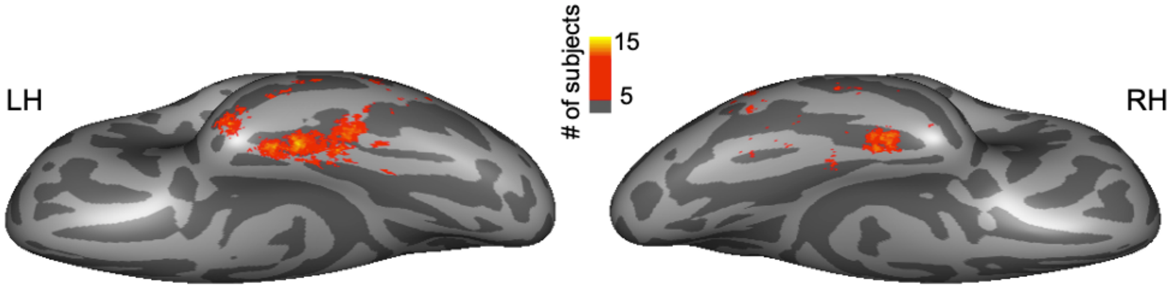


**Figure S6. Probabilistic maps for language and load-based attentional responses in the right VTC. Left**: the probabilistic map for Hard > Easy during the multiple-demand localizer (minimum overlap = 5 subjects) within the right VTC. **Right**: the probabilistic map for Sn>Tx showing subjects with overlapping activation during the language localizer (minimum overlap = 5 subjects) within the right VTC.

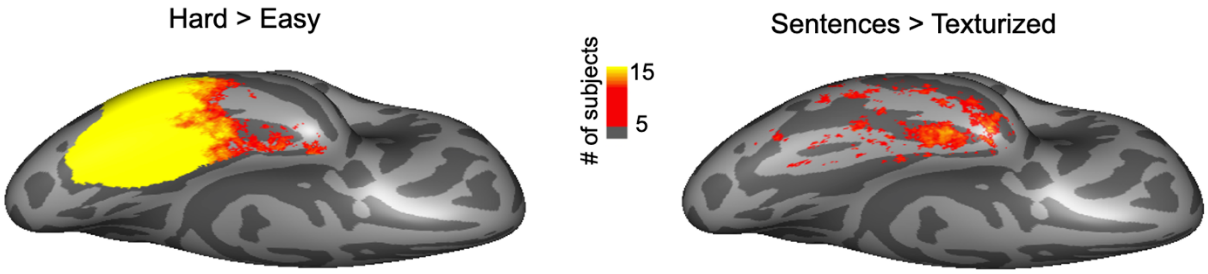

**Figure S7**: **Probabilistic maps for faces, objects, scenes, and bodies, created with the dynamic localizer**. For a given contrast, each subject's statistical map was thresholded at p<0.01 and the resulting binarized maps were added together. Here the probabilistic maps show vertices that overlap among at least 5 subjects.

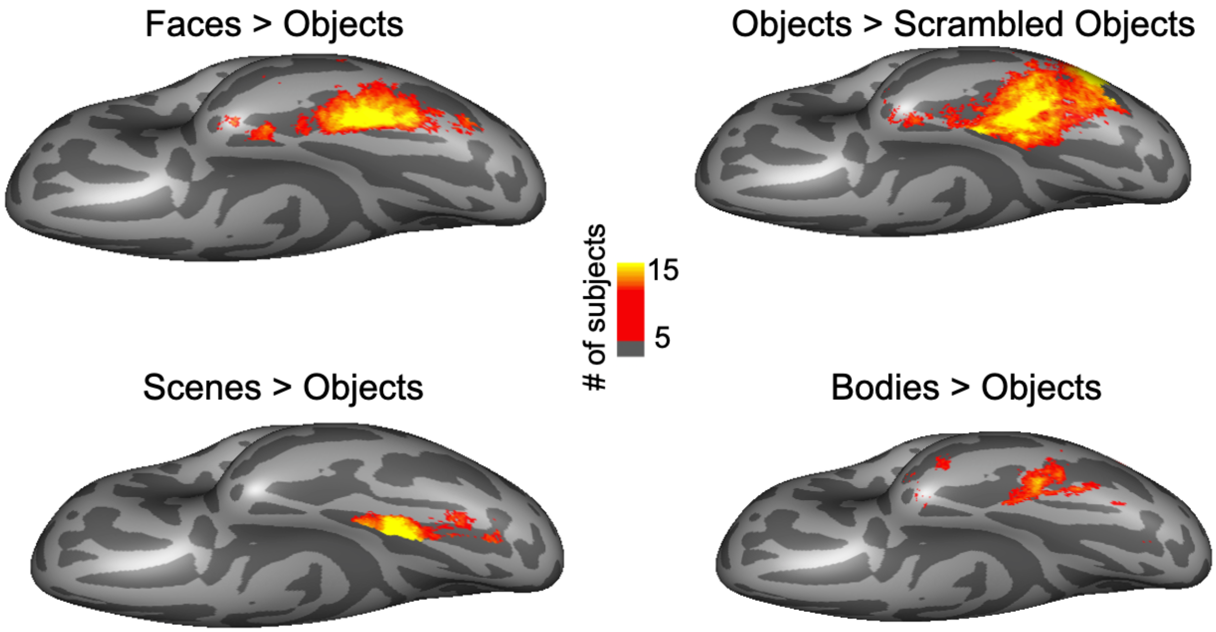

**Table S7: Descriptive information for fROIs when defined with a hard threshold and top 10%.**

Table S7a. Information for fROIs defined using a hard threshold ($p<0.005$).

| Number of subjects that have the fROIs (N Total = 37) | | | | | | | |
|---|---|---|---|---|---|---|---|
| Hemisphere | VWFA | FFA | OFA | FBA | PFS | PPA | RSC |
| Left | 35 | 36 | 24 | 34 | 35 | 37 | 37 |
| Right | 29 | 36 | 26 | 33 | 32 | 37 | 37 |
| Number of vertices in each ROI (Mean(SD)) | | | | | | | |
| Hemisphere | VWFA | FFA | OFA | FBA | PFS | PPA | RSC |
| Left | 222(208) | 191(159) | 43(67) | 130(116) | 433(339) | 892(458) | 944(408) |
| Right | 45(179) | 264(201) | 130(165) | 113(105) | 186(165) | 985(461) | 1050(481) |

Table S7b. Overlap between regions when defining the fROIs by selecting the top 10% vertices.

| **fROIs defined top 10% parcel** | | |
|---|---|---|
| **fROI 1 (mean size)** | **fROI 2 (mean size)** | **Overlapping vertices mean (sd)** |
| VWFA (597.27) | FFA (202.24) | 18 (20.25) |
| | FBA (175.76) | 10.83 (11.31) |
| | PFS (491.21) | 9.26 (12.59) |
| FFA (202.24) | FBA (175.76) | 4.41 (5.81) |
| | PFS (491.21) | 2.66 (6.61) |
| FBA (175.76) | PFS (491.21) | 5.85 (10.41) |
| PFS (491.21) | PPA (514.57) | 0.72 (2.50) |
